# TXNIP mediates LAT1/SLC7A5 endocytosis to reduce amino acid uptake in cells entering quiescence

**DOI:** 10.1101/2024.10.29.620655

**Authors:** Jennifer Kahlhofer, Nikolas Marchet, Brigitta Seifert, Kristian Zubak, Madlen Hotze, Anna-Sophia Egger, Claudia Manzl, Yannick Weyer, Sabine Weys, Martin Offterdinger, Sebastian Herzog, Veronika Reiterer, Marcel Kwiatkowski, Saskia B. Wortmann, Siamak Nemati, Johannes A. Mayr, Johannes Zschocke, Bernhard Radlinger, Kathrin Thedieck, Lukas A. Huber, Hesso Farhan, Mariana E.G. de Araujo, Susanne Kaser, Sabine Scholl-Bürgi, Daniela Karall, David Teis

## Abstract

Entry and exit from cellular quiescence require dynamic adjustments in nutrient acquisition, yet the mechanisms by which quiescent cells downregulate amino acid (AA) transport remain poorly understood. Here, we demonstrate that cells entering quiescence select plasma membrane-resident AA transporters for endocytosis and lysosomal degradation, to match AA uptake with reduced translation. We identify the α-arrestin TXNIP as a key regulator of AA uptake during quiescence, since it mediates the endocytosis of the SLC7A5-SLC3A2 (LAT1-4F2hc) transporter complex in response to reduced AKT signaling. Mechanistically, TXNIP interacts with HECT-type ubiquitin ligases to facilitate transporter ubiquitination. Loss of TXNIP disrupts this regulation, resulting in dysregulated AA uptake, sustained mTORC1 signaling, and accelerated quiescence exit. A novel TXNIP loss-of-function mutation in a patient with severe metabolic disease further supports its role in nutrient homeostasis and human health. These findings highlight TXNIP’s role in controlling SLC7A5-SLC3A2 mediated AA acquisition with implications for quiescence biology and disease.

## Introduction

How cells maintain a homeostatic pool of 20 proteinogenic amino acids (AA) is a fundamental question in biology. For the uptake and the release of AA across the plasma membrane (PM) and across organelle membranes, the human genome encodes at least 66 AA transporters, which belong to 11 individual solute carrier (SLC) families ^1,2^. The importance of controlling AA transport is exemplified by monogenic diseases and human pathologies caused by the deficiency or dysregulation of individual AA transporters ^3,4^.

Cellular AA import is coordinated with cell proliferation. Proliferating cells, in particular cancer cells, increase the abundance of AA transporters at the cell surface and thereby promote nutrient uptake for protein synthesis during cell growth prior to cell division ^5^. Transcriptional mechanisms are responsible for the upregulation of AA transporter expression ^6–9^.

Conversely, entry into quiescence, the reversible exit from the cell cycle, is accompanied by a reduction in cell size, lower rates of transcription, and reduced protein synthesis ^10,11^. Thus, quiescent cells must somehow re-calibrate AA uptake to align it with lower translation and metabolic homeostasis for cell survival, rather than for growth ^12^. How quiescent cells match AA flux across the PM to reduced cell size and to lower translation rates is unclear. This presents a knowledge gap in understanding cellular nutrient acquisition.

In budding yeast, *Saccharomyces cerevisiae*, we previously demonstrated that entry into quiescence induces the ubiquitin dependent endocytosis of AA transporters ^13^. The selective ubiquitination of AA transporters requires proteins of the α-arrestin family ^14^ ^15^. Activated α-arrestins function as adaptor proteins that recruit the homologous to the E6-AP carboxyl terminus (HECT) type ubiquitin ligase Rsp5 to different AA transporters for ubiquitination, endocytosis and lysosomal degradation ^16–18^.

The human genome encodes six α-arrestin family members or ARRDCs (arrestin-domain-containing proteins). With the exception of ARRDC5, α-arrestins contain one or more PPxY motifs in their C-terminal regions and are able to interact with several HECT-type ubiquitin ligases ^19–22^. The best characterized α-arrestin is thioredoxin interacting protein (TXNIP). TXNIP was first described as a negative regulator of thioredoxin (TRX) through the formation of an intermolecular disulfide bond that requires two critical cysteine residues of TXNIP (cysteine 32 and cysteine 247) ^23–25^.

TXNIP also regulates glucose uptake by mediating the selective endocytosis of SLC2A1 (GLUT1) and SLC2A4 (GLUT4), two major glucose transporters. TXNIP binds to these glucose transporters at the PM and interacts with a di-leucine motif in its C-terminal tail with components of the endocytic machinery ^26,27^. The TXNIP dependent regulation of SLC2A1 and SLC2A4 is tightly coupled to intracellular glucose homeostasis, ATP availability and growth factor signaling. AMPK activation in response to low ATP levels or AKT activation in response to insulin signal results in the phosphorylation of TXNIP at serine 308, thereby inactivating it ^26,27^. This inactivation of TXNIP ensures SLC2A1 and SLC2A4 accumulation at the PM to increase glucose uptake ^26,27^. TXNIP inactivation can involve ubiquitination by the HECT-type ubiquitin ligase ITCH, leading to its proteasomal degradation ^19,22^. These processes appear to be largely independent of TRX binding. Consistent with the function in regulating glucose uptake, biallelic loss-of-function mutations in TXNIP in humans lead to lactic acidosis in three siblings. Interesting, these patients also showed lower serum methionine levels ^28^.

Here, we show that in human cells entering quiescence, several AA transporters are selectively removed from the PM by endocytosis, including the heterodimeric AA transporter (HAT) SLC7A5-SLC3A2 (LAT1-4F2hc). The endocytic degradation of SLC7A5-SLC3A2 during entry into quiescence requires TXNIP. TXNIP uses its PPxY motifs to engage HECT-type ubiquitin ligases for SLC7A5-SLC3A2 endocytosis and subsequent lysosomal degradation. The TXNIP mediated endocytic degradation of SLC7A5-SLC3A2 reduces AA uptake, lowers free intracellular AA levels, and thereby helps to dampen mTORC1 signaling and translation. In growing cells, AKT phosphorylation of TXNIP restricts SLC7A5-SLC3A2 endocytosis. In addition, we identified a novel biallelic loss-of-function TXNIP variant in a boy with a severe inborn metabolic disease. The analysis of fibroblasts from this patient revealed that SLC7A5-SLC3A2 endocytosis was also blocked. This further supports the central role of TXNIP in adjusting cellular AA uptake, with important implications for human health.

## Results

### Selective endocytosis and lysosomal degradation of AA transporters in cells entering quiescence

To study the regulation of nutrient uptake during the transition from proliferation to quiescence, we used hTERT RPE1 cells (non-cancerous human telomerase-immortalized retinal pigmented epithelial, hereafter RPE1) as a model cell line ^29,30^. First, we established and evaluated conditions for entry into quiescence.

The DNA content of cells growing with serum was analyzed by propidium iodide (PI) staining and fluorescence activated cell sorting (FACS). 46% (± 2%) of these cells were in the G0/G1 phase of the cell cycle, 30% (± 6%) were in S phase and 24% (± 4%) in the G2/M phase. 24 h after serum starvation (but in the presence of AA and glucose), the number of cells in G0/G1 increased to 67% (± 10%), while the number of cells in S-phase and G2/M phase decreased to 15% (± 6%) and 19% (± 7%), respectively (Figure 1a). Cell viability was not affected by serum starvation (Supplementary Figure 1a). Serum starved cells also decreased their volume by 20% from 2907 fl to 2332 fl (Figure 1b). Consistent with an entry into quiescence in response to serum starvation, SDS-PAGE and Western Blot (WB) analysis of total cell lysates showed that the protein levels of the cyclin-dependent kinase inhibitor p27^Kip1^ increased, while the phosphorylation dependent activation of mitogen activated kinases ERK1/2, and mTORC2-dependent phosphorylation of AKT on serine 473 decreased (Figure 1c, lane 2).

**Figure 1.**
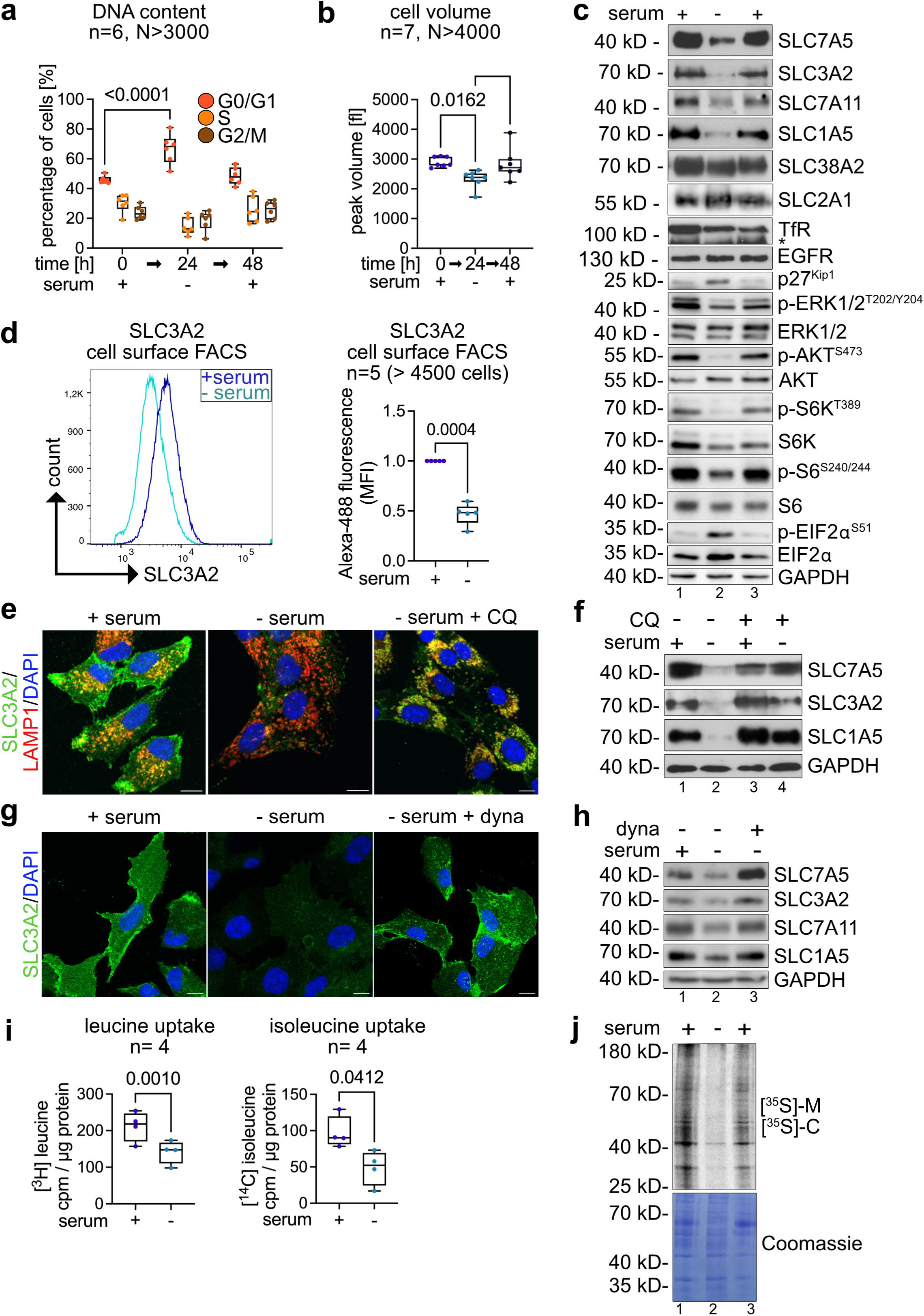
Dynamin-dependent endocytosis and lysosomal degradation of SLC7A5 and SLC1A5. RPE1 cells were grown in DMEM supplemented with serum (0 h, + serum), cultured in serum free medium for 24 h (24 h, -serum) and refed with serum for 24 h (48 h, + serum). **(a)** Cells were harvested, permeabilized and fixed, and the DNA content was analyzed by propidium iodide (PI) staining to assess the cell cycle profile by FACS (n=6, N>3000 cells, two-way ANOVA, Tukey’s multiple comparisons test). **(b)** Peak volume (in fl) was assessed by CASY Cell Counter and Analyzer (n=7, N>4000 cells, one-way ANOVA, Tukey’s multiple comparisons test). **(c)** Total cells lysates were analyzed by SDS-PAGE and Western Blot (WB) with the indicated antibodies. **(d)** The cell surface was stained with anti-SLC3A2 and anti-mouse Alexa-488. FACS was used to detect SLC3A2 surface staining. A representative histogram is shown. Quantification of the fluorescence intensities of SLC3A2 was normalized to proliferating cells (+ serum) (n=5, N>4500 cells, paired t-test). **(e, g)** Indirect immunofluorescence (IF) of PFA fixed cells stained for SLC3A2 (green), LAMP1 (red) and DAPI (blue) were analyzed by confocal microscopy. The merged images show a single plane of a Z-stack. Scale bar = 10 µm. **(e)** Incubation with 12.5 µM chloroquine (CQ) in absence of serum for 14 h (-serum, + CQ). **(g)** Incubation with 20 µM dynasore (dyna) in absence of serum for 14 h (-serum, + dyna). **(f, h)** Total cells lysates of cells treated as in (e, g) were analyzed by SDS-PAGE and WB with the indicated antibodies. **(i)** Cells were incubated with [^3^H]-leucine or [^14^C]-isoleucine for 15 min, washed and lysed. Total cell lysates were analyzed by scintillation counting. Counts per minute (cpm) values were normalized to total protein content (cpm / µg protein, n=4, paired t-test). **(j)** Cells were incubated with [^35^S]-methionine, [^35^S]-cysteine for 2 minutes, before cycloheximide (CHX, 10 µg/ml) was added, cells were lysed and analyzed by SDS-PAGE and autoradiography.

Of note, in cells entering quiescence, the protein levels of the heterodimeric AA transporters (HAT) SLC7A5-SLC3A2 (LAT1-4F2hc) and SLC7A11-SLC3A2 (xCT-4F2hc) decreased (Figure 1c, lane 1 and 2). Also, a decrease in the protein levels of SLC1A5 (ASCT2) was detected (Figure 1c, lane 1 and 2). The protein levels of the neutral AA transporter SLC38A2 (SNAT2), the glucose transporter SLC2A1 (GLUT1), the transferrin receptor TfR and epidermal growth factor receptor (EGFR) remained relatively unchanged over the course of the experiment (Figure 1c, lane 1 - 3). The downregulation of SLC7A5 and SLC1A5 was observed in other non-cancerous cell lines, such as mouse embryonic fibroblasts (MEFs) (Supplementary Figure 1b) and primary human fibroblasts (Supplementary Figure 1c, d), when subjected to serum starvation. However, this downregulation was not observed in a panel of seven different lung cancer cell lines (Supplementary Figure 1e).

To understand how SLC7A5-SLC3A2 and SLC1A5 were downregulated in cells entering quiescence, we examined the localization of these transporters using indirect immunofluorescence (IF) and confocal microscopy. In growing cells, endogenous SLC3A2 (Figure 1e) and SLC1A5 (Supplementary Figure 1f) were mainly detected at the PM and partially co-localized with LAMP1 positive lysosomes (Figure 1e and Supplementary Figure 1f). In serum starved cells, signals for SLC3A2 were barely detected (Figure 1e), which was in line with the downregulation of SLC7A5-SLC3A2 in quiescent cells. Also, cell surface FACS analysis measured a marked decrease of PM resident SLC3A2 (Figure 1d). Treatment of cells with chloroquine (CQ), which impairs lysosomal catabolic function, resulted in the accumulation of SLC3A2 (Figure 1e) and SLC1A5 (Supplementary Figure 1f) inside LAMP1 positive lysosomes and efficiently prevented the lysosomal degradation of these AA transporters (Figure 1f). Inhibition of the GTPase dynamin with dynasore (dyna), which blocks the scissions of clathrin coated vesicles from the PM ^31,32^, impaired the endocytosis of SLC3A2 (Fig 1g) and SLC1A5 (Supplementary Figure 1g) and in consequence prevented their lysosomal degradation (Figure 1h).

In line with the quiescence induced endocytic removal of SLC7A5-SLC3A2 from the PM, the uptake of its AA substrates [^3^H]-leucine and [^14^C]-isoleucine was reduced by 33% and 49%, respectively (Figure 1i). Similarly, the uptake of the SLC1A5 substrate [^14^C]-glutamine decreased by 20% (Supplementary Figure 1g).

The decrease in AA uptake upon entry into quiescence correlated with a reduction of mTORC1 signaling (*e.g.* towards S6K) and with an increase of eIF2α phosphorylation at serine 51 (Figure 1c, line 1 and 2), indicating a downregulation of translation. Consistently, the incorporation of [^35^S]-methionine and [^35^S]-cysteine into newly synthesized proteins was reduced in quiescent cells (Figure 1j, line 1 and 2).

Upon re-addition of serum to starved cells, SLC7A5-SLC3A2, SLC7A11-SLC3A2, and SLC1A5 protein levels were upregulated, ERK-, AKT-as well as mTORC1-signaling were re-activated, phosphorylation of eIF2α decreased as did p27^Kip1^ protein levels (Figure 1c, lane 3). Translation (Figure 1j, lane 3) and cell size increased (Figure 1c), and cells resumed proliferation (Figure 1a).

These results showed that the selective endocytosis and degradation of SLC7A5-SLC3A2 coincided with a decrease in cell volume, with a drop in translation, and with a reduction in (iso)leucine uptake as cells entered quiescence in response to serum starvation.

### TXNIP is required for the selective downregulation of SLC7A5

To identify the molecular mechanism responsible for the quiescence induced endocytic degradation of SLC7A5-SLC3A2, we focused on the α-arrestin family of ubiquitin ligase adaptors. α-arrestins are required for the endocytic downregulation of AA transporters in budding yeast cells entering quiescence ^13,33^. It was not known if they had a similar function in human cells.

The human genome encodes six α-arrestins, called ARRDC (arrestin domain containing) 1-5, and TXNIP ^15^. TXNIP protein levels (Figure 2a, b) and mRNA levels (Supplementary Figure 2a) increased in response to serum starvation.

**Figure 2.**
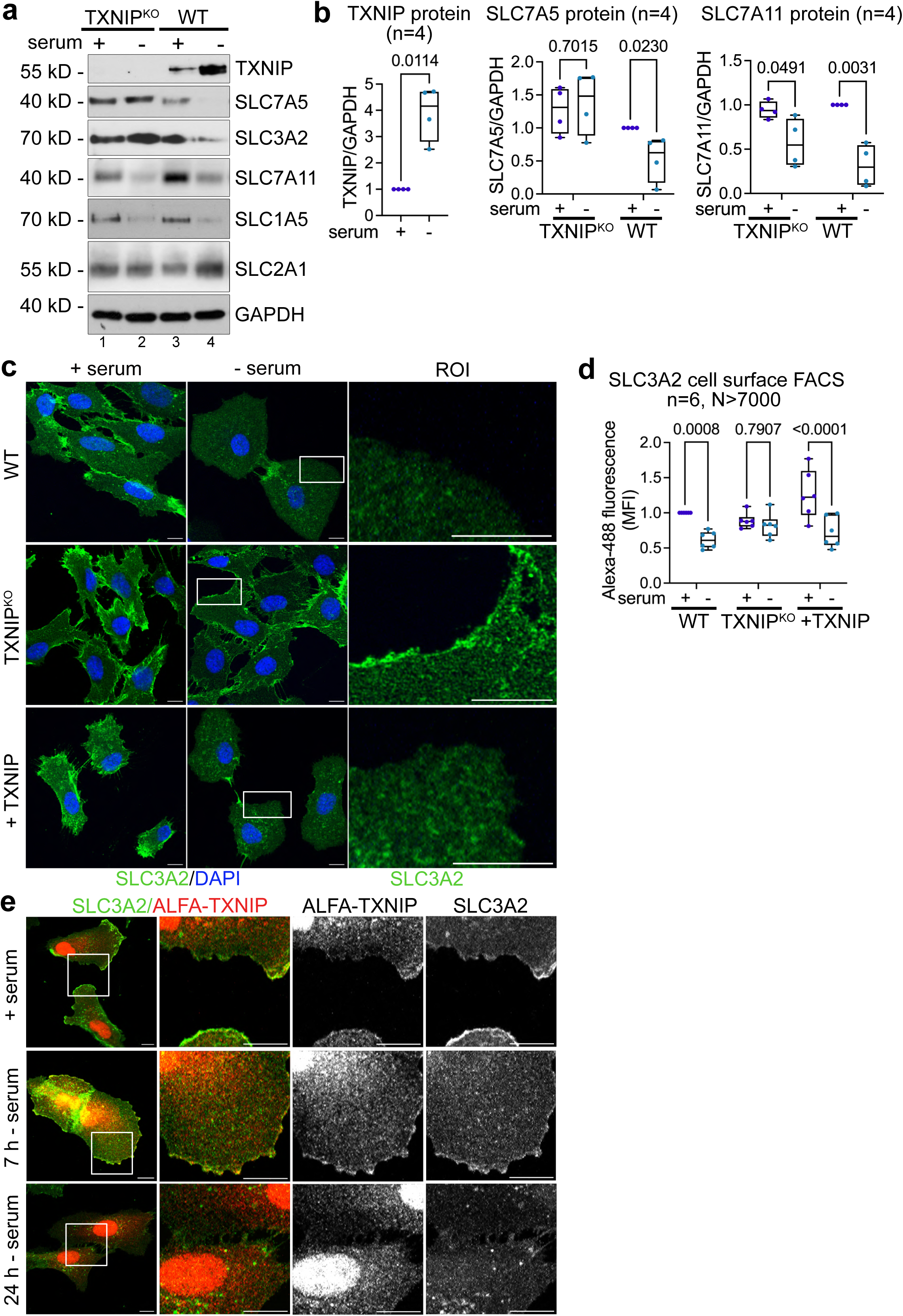
The importance of TXNIP for the selective degradation of SLC7A5. **(a)** TXNIP^KO^ and WT cells were grown in growth medium (+ serum) or serum starved for 24 h (-serum). Total cell lysates were analyzed by SDS-PAGE and WB with the indicated antibodies. **(b)** WB quantification of TXNIP (n=4, paired t-test), SLC7A5 (n=4, two-way ANOVA, Sidak’s multiple comparisons test) and SLC7A11 protein levels (n=4, two-way ANOVA, Sidak’s multiple comparisons test). Values were normalized to proliferating WT cells (+ serum). **(c)** IF of PFA fixed cells (WT, TXNIP^KO^ and TXNIP^KO^ reconstituted with TXNIP) stained for SLC3A2 (green) and DAPI (blue) were analyzed by confocal microscopy. The images show a single plane of a Z-stack. Regions of interest (ROI, white box) were magnified. Scale bar = 10 µm. **(d)** Quantification of SLC3A2 cell surface FACS of the indicated cells, normalized to proliferating WT cells (n=6, N>7000, two-way ANOVA, Sidak’s multiple comparisons test). **(e)** IF of PFA fixed TXNIP^KO^ reconstituted with ALFA-TXNIP. The localization of ALFA-TXNIP (red) and SLC3A2 (green) was analyzed by confocal microscopy. The images show a single plane of a Z-stack. Regions of interest (ROI, white box) were magnified. Scale bar = 10 µm.

To determine whether TXNIP contributed to SLC7A5-SLC3A2 endocytosis and lysosomal degradation, we used CRISPR/Cas9 mediated gene editing and introduced a 13 base pair deletion at position c.644_656 leading to a frameshift that resulted in a premature stop codon at position 226 (p.I215TfsTer11) (Supplementary Figure 2b, c). This mutation disrupted the architecture of the arrestin C-domain (Supplementary Figure 2b) and in single cell clones, the mutant protein was no longer detected (Figure 2a).

The loss-of-function mutation in TXNIP selectively prevented the endocytic downregulation of the SLC7A5-SLC3A2 heterodimer (Figure 2a, lane 2, and Figure 2b). In TXNIP^KO^ cells, SLC3A2 remained at the PM in response to serum starvation, as determined by IF and cell surface FACS (Figure 2c, d and Supplementary Figure 2d). Re-expression of TXNIP in TXNIP^KO^ cells restored SLC7A5-SLC3A2 endocytosis and degradation in response to serum starvation (Figure 2c, d and Supplementary Figure 2d - f). Conversely, TXNIP was not required for the degradation of SLC7A11 and SLC1A5 (Figure 2a, lane 2 and Supplementary Figure 2b). Hence, in cells entering quiescence, TXNIP selectively targeted SLC7A5-SLC3A2 for endocytosis and degradation. Loss of TXNIP did not affect SLC7A5 mRNA levels (Supplementary Figure 2g).

To examine the localization of TXNIP, ALFA-tagged TXNIP (ALFA-TXNIP) was re-expressed in TXNIP^KO^ cells. Using confocal microscopy, we detected a fraction of ALFA-tagged TXNIP at the PM, where it partially co-localized with SLC3A2 ^27,34^ in proliferating cells, and 7 h after serum starvation (Figure 2e). 24 h after serum starvation, the majority of SLC3A2 was endocytosed and also ALFA-TXNIP was no longer detected at the PM (Figure 2e). Under all conditions, ALFA-TXNIP also localized prominently to the nucleus and to the cytosol, as previously observed ^35^ (Figure 2e).

TXNIP has been shown to be required for controlling cellular glucose uptake, by regulating clathrin mediated endocytosis of the glucose transporters SLC2A1 and SLC2A4 in response to glucose depletion or insulin signaling ^26,27,36,37^. In WT and in TXNIP^KO^ RPE1 cells entering quiescence, SLC2A1 remained at the PM (Supplementary Figure 2h) and the protein levels of SLC2A1 rather increased in response to serum starvation (Supplementary Figure 2i). Consistently, the uptake of the fluorescent glucose analog 2-NBDG (2-[N-(7-nitrobenz-2-oxa-1,3-diazol-4-yl)amino]-2-deoxy-D-glucose) also increased in cell entering quiescence (Supplementary Figure 2j) ^38^. In the TXNIP^KO^ cells, 2-NBDG uptake was moderately elevated already in growing cells (Supplementary Figure 2j).

Our data suggested that TXNIP was required for the selective endocytic degradation of the heterodimer SLC7A5-SLC3A2 in cells entering quiescence. Since entering into quiescence did not trigger SLC2A1 endocytosis, it seemed that the TXNIP mediated downregulation of SLC7A5-SLC3A2 was not linked to TXNIP mediated SLC2A1 endocytosis.

### Identification of a novel TXNIP loss-of-function mutation in a patient with an inborn metabolic disorder

TXNIP deficiency caused by a loss-of-function variant c.174_175delinsTT has been previously reported in three sibling with congenital lactic acidosis, low serum methionine levels, variable hypoglycaemia and other metabolic alterations ^28^. During the course of the study, we identified a novel TXNIP loss-of-function mutation in another patient. The boy was born in 2014 and presented immediately after birth with recurrent hypoglycemia and muscular hypotonia in conjunction with lactic acidosis (Supplementary Table 1). Subsequently, he showed global developmental delay and marked intellectual disability with autistic features. In addition, he developed epileptic seizures with increasing frequencies of up to 3-4 per month (controlled with levetiracetame, topiramate and lamotrigine). With time, blood glucose concentrations stabilized (Supplementary Table 1), but the boy experienced recurrent hypoglycaemia in response to metabolic stress, e.g., infections. There were consistent metabolic alterations compatible with an AA transporter deficiency. Blood plasma concentrations of several large neutral amino acids (LNAAs, including L, I, V) were elevated throughout the years 2014 – 2022 (Supplementary Table 1). The increased molar ratio of the LNAAs (L, I, V) to aromatic AAs (F, Y), resulted in an elevated Fischer’s ratio (FR, 2014: FR = 4.46; 2016: FR = 5.38, 2018: FR = 5.90; 2021; FR=6.98; 2022: FR = 4.23; FR reference range = 2.10 -4). The methionine levels are not dramatically altered (Supplementary Table 1).

Whole-exome sequencing from peripheral blood of the patient detected a homozygous single nucleotide insertion c.642_643insT in exon 5 of 8 of the *TXNIP* gene, which was not recorded in the population genetic variant database gnomAD that lists *TXNIP* as likely haplosufficient (pLI = 0, LOEUF = 0,709: https://gnomad.broadinstitute.org accessed Sept. 10, 2024). The variant caused a frameshift and a premature stop codon after 59 AA (denoted p.Ile215TyrfsTer59), likely causing nonsense-mediated decay (NMD) or the synthesis of a severely truncated TXNIP protein (Figure 3a). Both parents are healthy heterozygous carriers for the variant. No other (likely) pathogenic variant was identified as explanation of the clinical features in the child.

**Figure 3.**
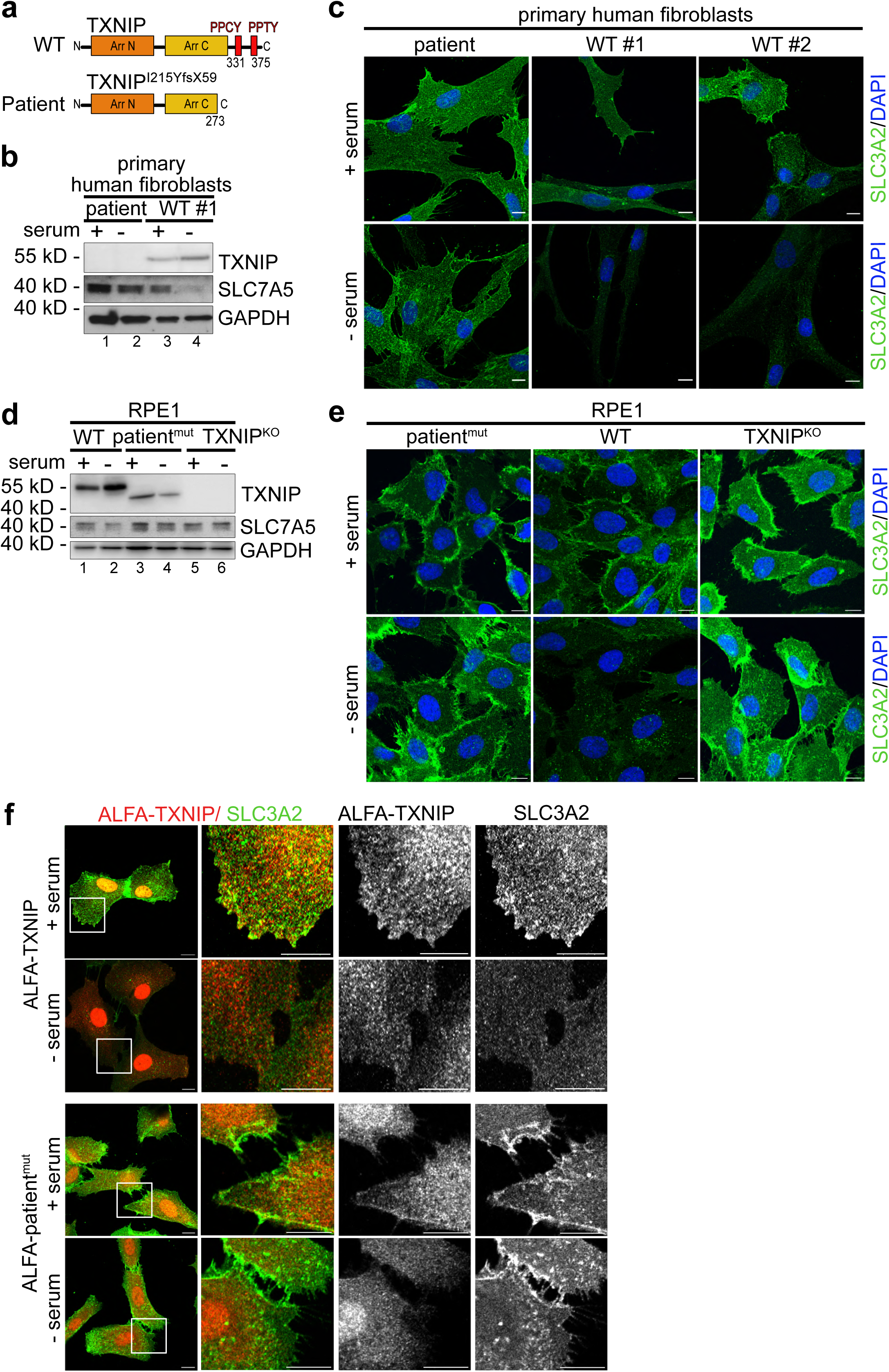
Loss of function mutation in TXNIP causes a rare disease. **(a)** Schematic representation of WT TXNIP and mutant TXNIP harboring the patient mutation (c.642_643insT, patient^mut^). **(b, c)** Control fibroblasts (WT#1 and WT#2) and patient-derived fibroblasts (patient) were grown in growth medium (+ serum) or serum starved for 24 h (- serum). **(b)** Total cell lysates were analyzed by SDS-PAGE and WB with the indicated antibodies. **(c)** Indirect IF of PFA fixed fibroblasts (patient, WT#1, WT#2) stained for SLC3A2 (green) and DAPI (blue) was analyzed by confocal microscopy. The images show a single plane of a Z-stack. Scale bar = 10 µm. (**d, e)** RPE1 WT, TXNIP^KO^ and TXNIP^KO^ reconstituted with ALFA-TXNIP harboring the patient mutation (c.642_643insT, patient^mut^) were grown in growth medium (+ serum) or serum starved for 24 h (- serum). **(d)** Total cell lysates were analyzed by SDS-PAGE and WB with the indicated antibodies. **(e)** Indirect IF of PFA fixed cells (WT, TXNIP^KO^ and patient^mut^) stained for SLC3A2 (green) and DAPI (blue) was analyzed by confocal microscopy. The images show a single plane of a Z-stack. Scale bar = 10 µm. **(f)** ALFA-TXNIP (red) and SLC3A2 (green) was analyzed by confocal microscopy. The images show a single plane of a Z-stack. Regions of interest (ROI, white box) were magnified. Scale bar = 10 µm.

To examine how the loss-of-function mutation in TXNIP affected SLC7A5 downregulation, we generated primary patient-derived fibroblasts. WB analysis of the patient fibroblasts showed complete loss of the TXNIP protein (Figure 3b). In these cells, SLC7A5 was no longer downregulated in response to serum starvation (Figure 3b). Furthermore, indirect IF and confocal microscopy showed that SLC3A2 remained at the PM in response to serum starvation in the patient-derived fibroblasts with the TXNIP loss-of-function mutation, but not in the control primary fibroblasts (Figure 3c).

To directly test if the patient mutation was causative for the failure to downregulate SLC7A5-SLC3A2, we expressed an ALFA-tagged version cDNA encoding TXNIP p.Ile215TyrfsTer59 (ALFA-patient^mut^, or patient^mut^) in RPE1 TXNIP^KO^ cells, under the control of a constitutive eIF1α promoter. Under these conditions (which do not trigger NMD), the expression of a truncated TXNIP protein was detected (Figure 3d, lane 3 and 4). ALFA-patient^mut^ localized to the PM, but failed to mediate SLC7A5-SLC3A2 endocytosis in response to serum starvation (Figure 3d-f). Hence, even if the protein was expressed, the TXNIP patient^mut^ protein was non-functional.

It seemed that a loss-of-function mutation in TXNIP caused a severe metabolic disease, inhibited the downregulation of SLC7A5-SLC3A2 in patient-derived fibroblasts upon entry into quiescence and caused persistent changes in AA levels in the serum of the affected patient.

### PPxY motifs of TXNIP were required for the endocytosis of SLC7A5

TXNIP contains two PPxY motifs in its C-terminal region (Figure 4a), which enable the interaction with different HECT type ubiquitin ligases that contain a WW domain ^15,19–22,39,40^. To test how the PPxY motifs of TXNIP contributed to SLC7A5-SLC3A2 endocytosis, we introduced point mutations either in both PPxY motifs (TXNIP^PPCY331AACA, PPTY375AATA^, hereafter TXNIP^PPxYmut^), or only in the first PPxY motif (TXNIP^PPCY331AACA^, hereafter TXNIP^PPxY331^) and expressed these mutants in TXNIP^KO^ RPE1 cells. The protein levels of the PPxY mutants were higher compared to endogenous TXNIP (Figure 4b and Supplementary Figure 3a), yet the endocytic downregulation of SLC7A5-SLC3A2 was blocked (Figure 4b and Supplementary Figure 3a) and SLC3A2 remained at the PM in response to serum starvation (Figure 4c, d and Supplementary Figure 3a -c).

**Figure 4.**
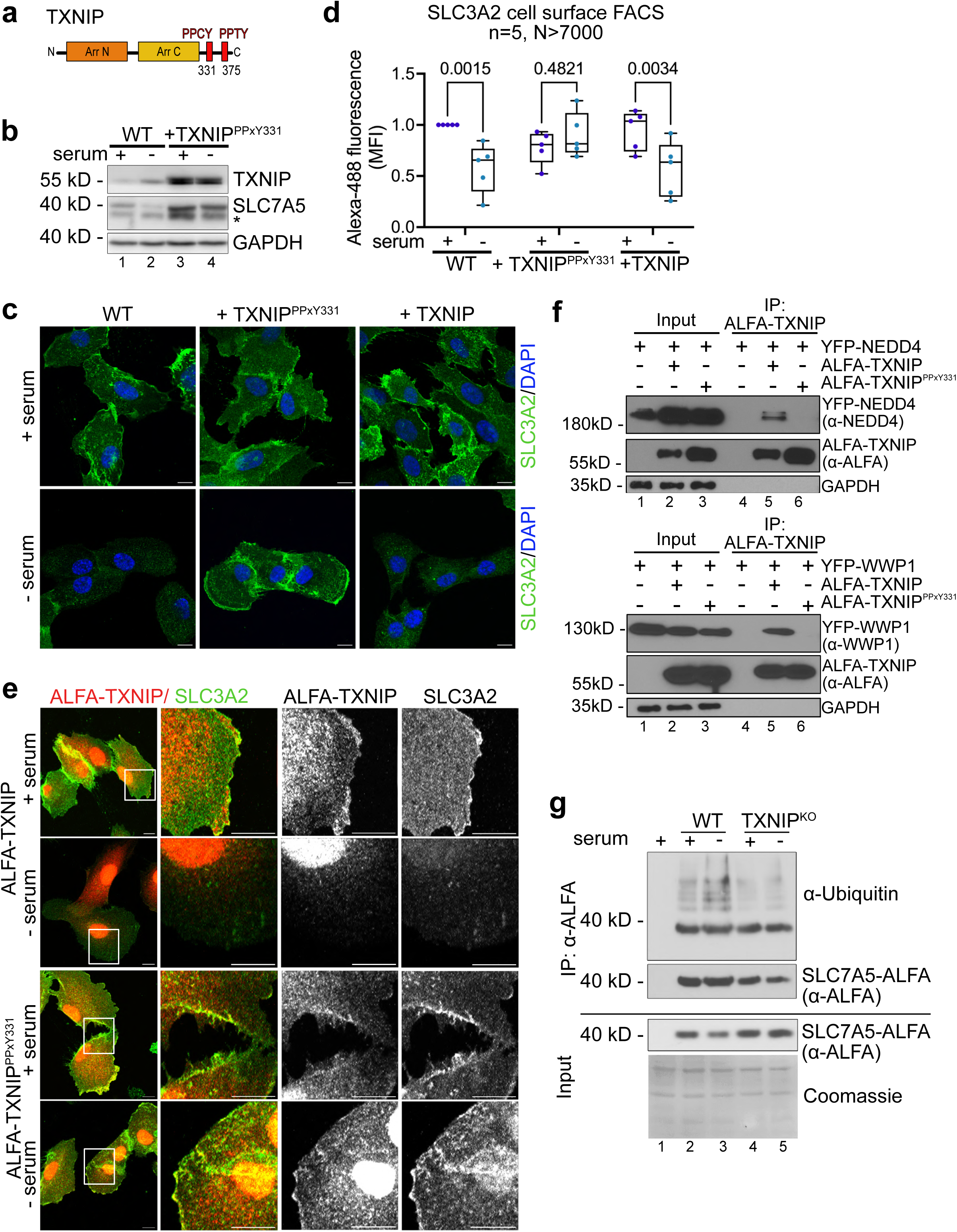
PPxY motifs of TXNIP in the downregulation of SLC7A5. **(a)** Schematic representation of TXNIP protein harboring PPxY motifs. **(b)** Total cell lysates from WT cells or TXNIP^KO^ reconstituted with ALFA-TXNIP^PPxY331^ mutants were analyzed by SDS-PAGE and WB with the indicated antibodies. The asterisk (*) labels a second band that was only detected using the monoclonal SLC7A5 antibody. **(c, e)** Indirect IF of PFA fixed cells (WT, TXNIP^KO^ and TXNIP^KO^ reconstituted with ALFA-TXNIP or ALFA-TXNIP^PPXY331^) was analyzed by confocal microscopy. The images show a single plane of a Z-stack **(c)** Cells were stained for SLC3A2 (green) and DAPI (blue). Scale bar = 10 µm. **(d)** Quantification of SLC3A2 cell surface FACS of the indicated cells, normalized to proliferating WT cells (n=5, N>7000 cells, two-way ANOVA, Sidak’s multiple comparisons test). **(e)** Cells were stained for ALFA-TXNIP (red) and SLC3A2 (green). Regions of interest (ROI, white box) were magnified. Scale bar = 10 µm**. (f)** SDS-PAGE and WB analysis with the indicated antibodies from non-denaturing ALFA-immunoprecipitations (IP) from HEK293T cells expressing ALFA-TXNIP or the mutated ALFA-TXNIP^PPxY331^ variant and the indicated YFP-tagged HECT-type ubiquitin ligase. **(g)** SDS-PAGE and WB analysis with the indicated antibodies from denaturing ALFA-SLC7A5 IP from SLC7A5^KO^ RPE1 reconstituted with ALFA-SLC7A5.

A fraction of ALFA-TXNIP^PPxY331^ localized to the PM in proliferating cells, similar to wildtype ALFA-TXNIP. In response to serum starvation, the signal for ALFA-TXNIP^PPxY331^ persisted at the PM, where it co-localized with SLC3A2, while ALFA-TXNIP was cleared from the PM together with SLC3A2 (Figure 4e). These results indicated that the endocytic downregulation of SLC7A5-SLC3A2 required the PPCY motif at position 331 of TXNIP.

To examine the capacity of TXNIP and its PPxY motif to interact with different HECT-type ubiquitin ligases, we co-expressed ALFA-TXNIP or ALFA-TXNIP^PPxY331^ with YFP-NEDD4, -ITCH, -WWP1, WWP2, and -HECW1 ^41, 42^ in HEK293T cells. A robust interaction of ALFA-TXNIP with YFP-NEDD4, YFP-WWP1 and YFP-WWP2 was detected (Figure 4f and Supplementary Figure 3d), which required the PPCY motif at position 331 in TXNIP (Figure 4f and Supplementary Figure 3d). The interaction of TXNIP appeared to be weaker with HECW1, and the interaction with ITCH was barely detected (Supplementary Figure 3d). It seemed that the PPCY motif at position 331 rendered TXNIP capable to interact with different HECT type ubiquitin ligases.

To examine if entry into quiescence induced TXNIP mediated ubiquitination of SLC7A5, we replaced endogenous SLC7A5 with C-terminally ALFA-tagged SLC7A5 (SLC7A5-ALFA). Therefore, we generated SLC7A5^KO^ cells (using CRISPR/Cas9 mediated gene editing, Supplementary Figure 3e), as well as SLC7A5^KO^ TXNIP^KO^ double knock out cells and stably over-expressed SLC7A5-ALFA. To detect ubiquitination, SLC7A5-ALFA was immunoprecipitated under denaturing conditions (1% SDS) before and after 7 h of serum starvation. Polyubiquitination of SLC7A5-ALFA was detected under steady state conditions, and appeared to slightly increase 7 h after serum starvation. The polyubiquitination of SLC7A5-ALFA was markedly reduced in the TXNIP^KO^ cells (Figure 4g). Hence, TXNIP seemed to be required for the efficient polyubiquitination of SLC7A5-ALFA.

Taken together, these findings suggested that TXNIP used its PPxY motif to interact with HECT-type ubiquitin ligases for the efficient ubiquitination of SLC7A5-SLC3A2, leading to endocytic downregulation.

### AKT signaling controls TXNIP mediated downregulation of SLC7A5

Earlier work demonstrated that AKT directly phosphorylated TXNIP on serine 308 to inhibit endocytosis of SLC2A1 and increase glucose uptake ^26^. To examine if AKT also controlled the endocytic downregulation of SLC7A5-SLC3A2, we treated RPE1 cells with the AKT inhibitor MK2206 (MK). MK2206 treatment caused the endocytic downregulation of the SLC7A5-SLC3A2 heterodimer (Figure 5a, lane 3 and Supplementary Figure 4a), but not of SLC1A5 (Supplementary Figure 4a). Inhibition of MEK with PD0325901 (PD) triggered the selective endocytic downregulation of SLC1A5, but not of SLC7A5-SLC3A2 (Figure 5a, lane 4 and Supplementary Figure 4a). These results suggested that distinct mitogenic signaling cascades regulated the PM levels of different AA transporters, with AKT controlling SLC7A5-SLC3A2 endocytosis.

**Figure 5.**
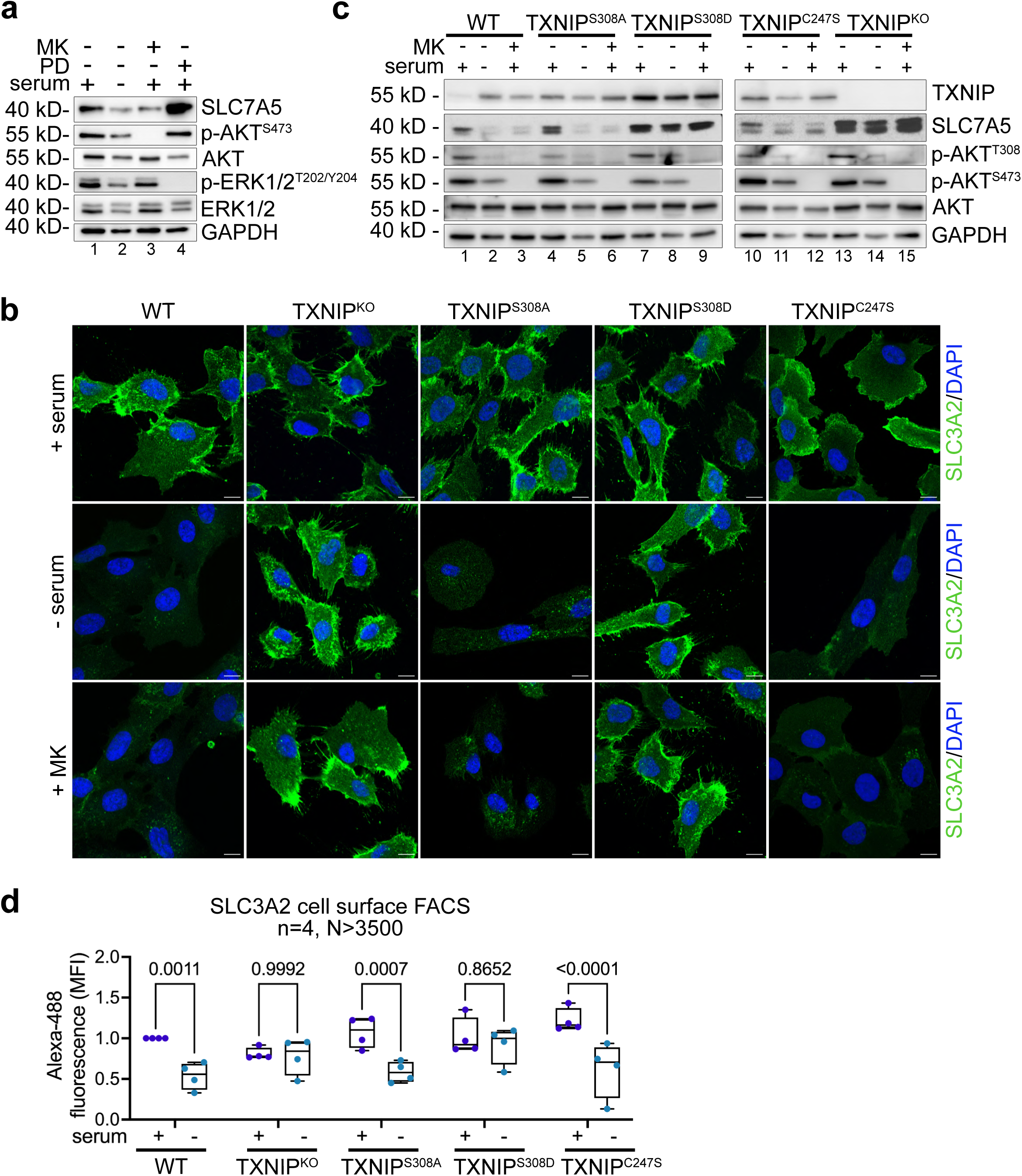
TXNIP and AKT signaling. **(a)** Total cell lysates from WT cells before or after 24 h of serum starvation or incubation with 1 µM MK2206 (MK) or 5 µM PD0325901 (PD). Cells were analyzed by SDS-PAGE and WB with the indicated antibodies. **(b)** Indirect IF of PFA fixed cells (WT, TXNIP^KO^ and TXNIP^KO^ reconstituted with TXNIP^S308A^, TXNIP^S308D^ or TXNIP^C247S^) before or after serum starvation or incubation with 1 µM MK2206 (MK) stained for SLC3A2 (green) and DAPI (blue). Cells were analyzed by confocal microscopy. The images show a single plane of a Z-stack. Scale bar = 10 µm. **(c)** Total cell lysates from cells treated as in (b) were analyzed by SDS-PAGE and WB with the indicated antibodies. **(d)** Quantification of SLC3A2 cell surface FACS from the indicated cells (n=4, N>3500 cells, two-way ANOVA, Sidak’s multiple comparisons test).

To test if AKT controlled SLC7A5 endocytosis via TXNIP, we treated RPE1 WT and TXNIP^KO^ cells with MK2206. In the TXNIP^KO^ cells, endocytosis and degradation of SLC3A2 was blocked in response to MK2206 treatment, suggesting that TXNIP functioned downstream of AKT to mediate SLC7A5-SLC3A2 endocytic degradation (Figure 5b, c, lane 1 – 3 and 13 - 15). Consistently, the phospho-mimetic TXNIP^S308D^ mutant blocked SLC3A2 endocytosis in response to serum starvation or AKT inhibition, while the non-phosphorylatable TXNIP^S308A^ mutant allowed SLC7A5-SLC3A2 endocytic degradation (Figure 5b, c). These findings were corroborated by cell surface FACS analysis of SLC3A2 (Figure 5d and Supplementary Figure 4b).

TXNIP forms a disulfide bound (S-S) via cysteine 247 to inhibit TRX (thioredoxin) activity ^23,24^. To test the role of the S-S between TXNIP and TRX in SLC7A5-SLC3A2 endocytosis, we generated mutant TXNIP^C247S^, which no longer formed a S-S bridge with TRX ^43^. The expression of TXNIP^C247S^ in TXNIP^KO^ cells rescued the endocytic degradation of SLC7A5-SLC3A2, similar to WT TXNIP (Figure 5b -d). Thus, SLC7A5 endocytosis was independent of the ability of TXNIP to form disulfide bridges with TRX.

These results demonstrated that AKT mediated phosphorylation of TXNIP at serine 308 inhibited SLC7A5-SLC3A2 endocytosis, while reduction of AKT signaling enabled TXNIP mediated SLC7A5-SLC3A2 endocytic degradation.

### TXNIP mediated downregulation of SLC7A5 was essential to adjust intracellular AA levels and metabolic signaling in cells entering quiescence

Our results so far linked growth factor signaling via AKT with the TXNIP mediated control of AA acquisition via SLC7A5-SLC3A2. To directly test whether TXNIP mediated endocytosis of SLC7A5-SLC3A2 adjusted AA uptake in cells entering quiescence, we measured the cellular uptake of [^3^H]-leucine. In response to serum starvation, WT cells downregulated [^3^H]-leucine uptake, while TXNIP^KO^ cells did not reduce [^3^H]-leucine uptake (Figure 6a). Upon JPH203 treatment, a selective inhibitor of SLC7A5 ^44^, the uptake of [^3^H]-leucine was blocked in WT cells and TXNIP^KO^ cells (Figure 6a), suggesting that SLC7A5 was the main transporter for leucine in RPE1 cells. Consistently, in SLC7A5^KO^ cells, the uptake of [^3^H]-leucine was strongly impaired (Supplementary Figure 5a).

**Figure 6.**
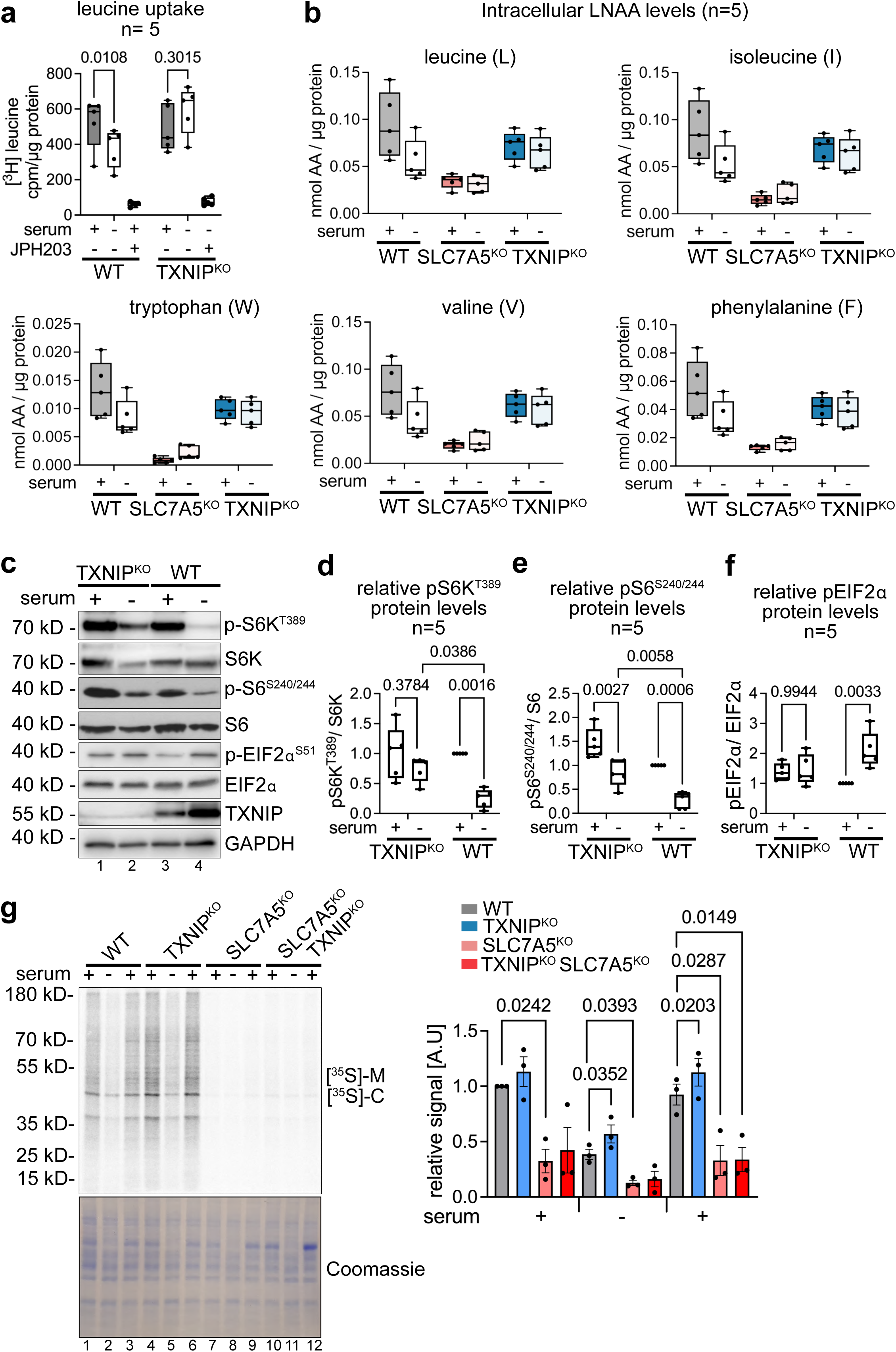
TXNIP mediated degradation of SLC7A5 contributed to metabolic signaling. **(a)** WT and TXNIP^KO^ cells were grown in growth medium (+ serum), serum starved for 24 h (- serum) or treated with 10 µM JPH203. Cells were incubated with [^3^H]-leucine for 15 min, washed and lysed. Cell lysates were analyzed by scintillation counting. Cpm values were normalized to total protein content (n=5, two-way ANOVA, Tukey’s multiple comparisons test). **(b)** WT, SLC7A5^KO^ and TXNIP^KO^ cells were grown in growth medium (+ serum) or serum starved for 24 h (- serum). Mass spectrometry analysis of free AAs, normalized to total protein content (nmol AA / µg protein, n=5). **(c)** Total cell lysates from WT and TXNIP^KO^ cells before or after 24h serum starvation, were analyzed by SDS-PAGE and WB with the indicated antibodies **(d-f)** WB quantification of pS6K^T389^, normalized to total S6K. WB quantification of pS6^S240/244^, normalized to total S6 and WB quantification of pEIF2α^S51^, normalized to total EIF2α protein levels (n=5, two-way ANOVA). **(g)** Cells were incubated with [^35^S]-methionine, [^35^S]-cysteine for 2 minutes, before CHX (10µg/ml) was added, cells were lysed and analyzed by SDS-PAGE and autoradiography follow by densitometric analysis (n=3, paired t-test).

To assess the effect of SLC7A5 downregulation or the lack thereof on intracellular free AA levels, we used quantitative mass spectrometry. We quantified free levels of 17 different proteinogenic AA in WT, TXNIP^KO^ and SLC7A5^KO^ RPE1 cells. In SLC7A5^KO^ cells the intracellular levels of L, I, W, V, F, Y were low under all conditions (Figure 6b), with exception for M, which was upregulated (Supplementary Figure 5b). These LNAAs are known substrates of SLC7A5 ^1^ (Figure 6b). In WT cells, the free levels of most these AAs (L, I, W, V, F, Y) decreased (Figure 6b). In contrast, the levels of L, I, W, V, F, Y were lower in growing TXNIP^KO^ cells, and they did not decrease when TXNIP^KO^ cells entered quiescence (Figure 6b). Neither loss of SLC7A5 nor loss of TXNIP affected the levels K, R, S, E, Q, P, G, A (Supplementary Figure 5b, c). Taken together, these data indicated that the loss of TXNIP affected the ability of cells to control SLC7A5 dependent AA transport.

To characterize the consequence resulting from the failure to adjust AA uptake in TXNIP^KO^ cells, we determined how mTORC1 dependent translational control was affected. In quiescent WT cells, phosphorylation of S6K on threonine 389 was reduced compared to proliferating cells, as was phosphorylation of the S6K substrate S6 on serine 240/244, in keeping with lower mTORC1 activity (Figure 6c, lane 3 and 4, Figure 6d, e). In TXNIP^KO^ cells, S6K and S6 phosphorylation also decreased, but remained higher compared to WT cells (Fig 6c, lane 1 and 2, Figure 6d, e), indicating elevated mTORC1 activity. Another important mode for translational control is eIF2α phosphorylation at serine 51 by GCN2 (but also other kinases of the integrated stress response) in response to lower AA levels. In quiescent WT cells, eIF2α phosphorylation at serine 51 increased (Figure 6c, lane 3 and 4, Figure 6f). However, in growing TXNIP^KO^ cells, eIF2α phosphorylation appeared to be slightly elevated but did not further increase, as these cells transitioned into quiescence (Figure 6c, lane 1 and 2, Figure 6f). It seemed that TXNIP mediated SLC7A5-SLC3A2 endocytosis was required to decrease AA uptake in quiescence, which helped to change signaling towards the translation machinery, possibly to adapt translation.

In line with this concept, [^35^S]-methionine and [^35^S]-cysteine incorporation into newly synthesized proteins remained higher in quiescent TXNIP^KO^ cells, and also appeared to increase faster after the re-addition of serum (Figure 6d). The increased translation in the TXNIP^KO^ cells required SLC7A5, because it was no longer observed in TXNIP^KO^ SLC7A5^KO^ double mutant cells (Figure 6g).

We concluded that TXNIP mediated endocytosis of SLC7A5-SLC3A2 reduced the uptake of LNAAs such as L, I, W, V, F, Y, which in turn lowered the free intracellular levels of these AA, thereby helping cells to reduce mTORC1 signaling and to dampen translational activity.

### TXNIP mediated endocytic degradation of SLC7A5 restrains cell proliferation

Finally, we determined how the TXNIP mediated regulation of SLC7A5 and AA uptake contributed to cell cycle progression during entry into and exit from quiescence. Therefore, we compared the DNA content of WT, TXNIP^KO^ and TXNIP^KO^ cells which re-expressed TXNIP during entry and exit from serum starvation induced quiescence (Figure 7a).

**Figure 7.**
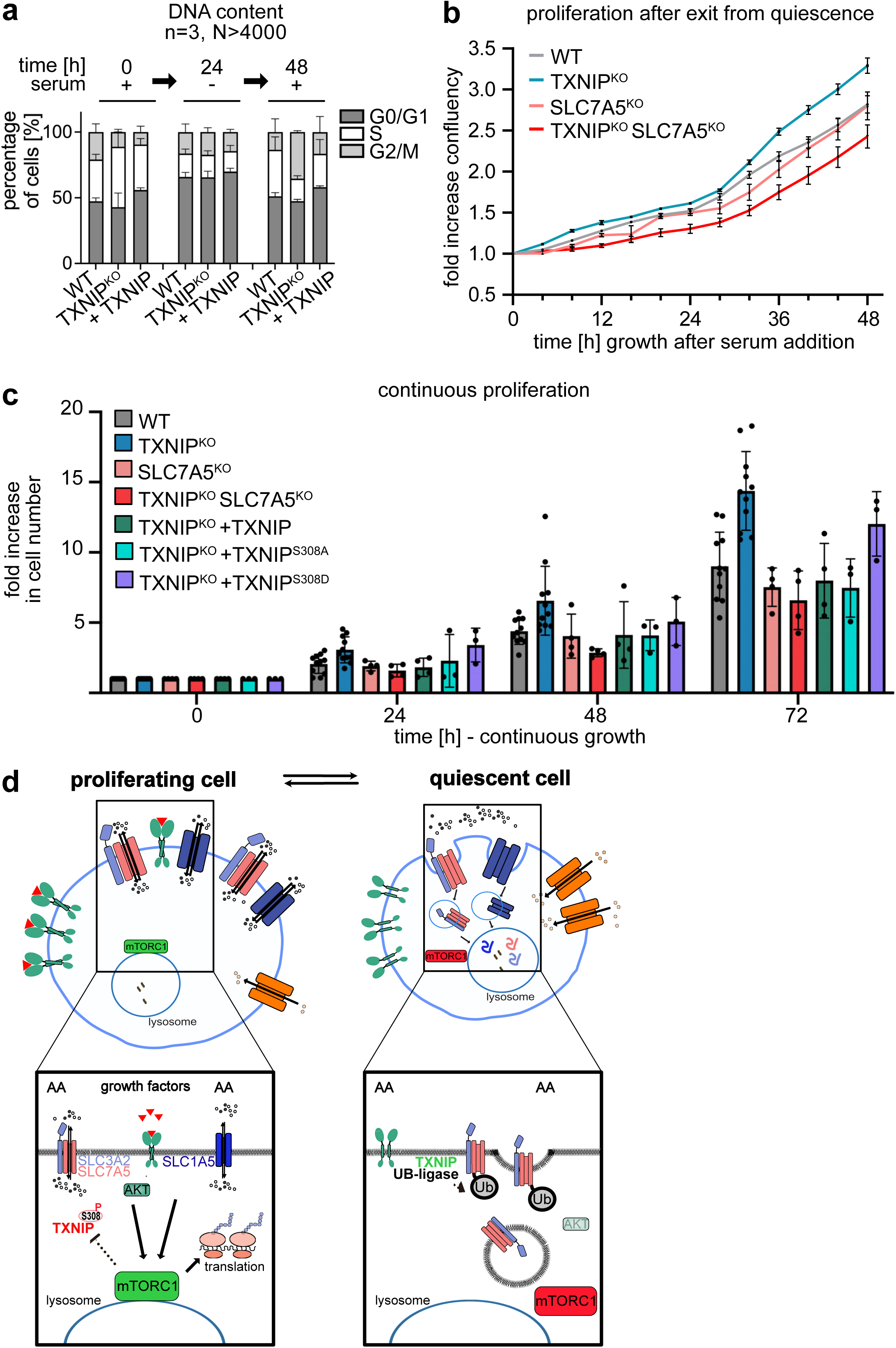
TXNIP mediated degradation of SLC7A5 contributed to the regulation of cell proliferation. **(a)** WT, TXNIP^KO^ and TXNIP^KO^ cells reconstituted with TXNIP were harvested, permeabilized and fixed. The DNA content was analyzed by PI staining to assess the cell cycle profile by FACS (n=3, N>4000 cells). **(b)** Increase in confluency of the indicated cells, after refeeding serum starved cells, was measured by continuous live cell microcopy in an Incucyte incubator for 48 h (n=6). **(c)** The proliferation of the indicated cells was determined using CASY cell counting during the indicated time. The relative increase in cell number is normalized to 0 h and show as fold increase (n>3). **(d)** Model of TXNIP mediated downregulation of SLC7A5 during entry into quiescence.

In a growing cell population and 24 h after serum starvation, the DNA content of WT, TXNIP^KO^ and rescued cells was similar (Figure 7a, timepoint 0 h and 24 h). Interestingly, 24 h after serum re-addition (timepoint 48 h), a larger fraction of the TXNIP^KO^ cells (35%, ± 1%) had already reached the G2/M phase of the cell cycle. In comparison, only 13% (± 10%) of WT cells or 17% (± 12%) of TXNIP^KO^ cells re-expressing TXNIP reached G2/M phase and more cells (35% ± 10% of WT cell and 25% ± 11% of rescued TXNIP^KO^) were still in S-Phase (Figure 7a). Consistent with a faster cell cycle progression of TXNIP^KO^ cells during exit from quiescence, these cells resumed cell proliferation faster compared to WT cells, which was dependent on SLC7A5 (Figure 7b). Moreover, the overall cell proliferation rate of TXNIP^KO^ cells or of TXNIP^KO^ cells expressing TXNIP^S308D^ was increased compared to WT or TXNIP^KO^ cells expressing TXNIP or TXNIP^S308A^ (Figure 7c). Again, the increased proliferation of TXNIP^KO^ cells was dependent on SLC7A5, since the double knock out TXNIP^KO^ SLC7A5^KO^ grew slower, similar to SLC7A5^KO^ cells (Figure 7c).

Taken together, our results demonstrated that TXNIP interacted with HECT-type ubiquitin ligases to selectively degrade SLC7A5-SLC3A2 in cells entering quiescence, thereby reducing AA uptake. This process not only contributed to reduce translation, but also restrained cell cycle progression during the exit from quiescence and limited overall cell proliferation (see Figure 7d model). This mechanism was only licensed upon serum removal and repressed by growth factor signaling through AKT.

## Discussion

Our work establishes a central role for TXNIP in a functional feedback loop that links growth factor signaling, AA homeostasis and metabolic signaling for translational control during cellular transitions between proliferation and quiescence, with potential relevance for human health. In particular, we made progress toward understanding how growth signaling regulates TXNIP mediated endocytosis of SLC7A5-SLC3A2 and hence, the acquisition of LNAAs for metabolic scaling in growing and quiescent cells. The framework of our model (Figure 7d) is based on the following results: (1) Human cells that enter quiescence downregulate AA uptake, in part, by selective endocytosis of the AA transporters; (2) the α-arrestin TXNIP mediates the selective endocytosis of SLC7A5-SLC3A2; (3) TXNIP mediated endocytosis is suppressed in growing cells by AKT signaling downstream of growth factor receptors and (4) TXNIP mediated SLC7A5-SLC3A2 endocytosis helps to lower AA uptake, translation and thereby restrains cell proliferation. (5) These processes contribute to metabolic homeostasis in humans. A novel loss-of-function mutation in TXNIP, that impairs AA endocytosis, causes a rare metabolic disease.

Together with a previous report on three siblings from a single consanguineous family that were homozygous for a loss-of-function frameshift variant c.174_175delinsTT ^28^, our data suggest TXNIP deficiency as a novel inherited metabolic disease with specific metabolic alterations, that appear associated with neonatal lactic acidosis, recurrent hypoglycaemia, and specific anomalies in AA metabolism, although full the spectrum of associated clinical manifestations remains uncertain. Similar to our patient, the three siblings presented with lactic acidosis in the neonatal period, which was treated with long-term dichloroacetic acid (DCA). Rather than directly clearing lactate from serum, DCA inhibits pyruvate dehydrogenase kinase to stimulate pyruvate dehydrogenase activity and oxidative decarboxylation of pyruvate, resulting in reduced serum lactate levels ^45^ ^46^. Two of the three affected children had normal development during childhood; one child, which also suffered from recurrent hypoglycaemia, had neonatal muscular hypotonia and failure to thrive, and was under investigation for autism spectrum disorder, similar to our patient. Our patient additionally has an increasing frequency of epileptic seizures. The three siblings developed low plasma methionine concentrations. No other AA anomalies were specified in the previous report except for a transient general hyperaminoaciduria in one child ^28^. The detailed metabolic investigations in our case revealed relative normal methionine levels, but identified other specific AA alterations in line with impaired SLC7A5 endocytosis in the case of complete TXNIP loss. How these metabolic defects are linked to other clinical manifestations such as possible autism and the severe epileptic developmental disorder in our patient, remains to be investigated. We speculate that these feature could be related to role of SLC7A5 in neurons and in the blood brain barrier ^47,48^. Yet, the patient phenotype appears to be variable in severity and not fully penetrant. Hence, loss of TXNIP alone is likely to be insufficient and other genetic or non-genetic factors (e.g. diet) that are not yet known, must be involved.

Several studies in mice show that TXNIP is important during the transition between fed and fasted states, which was attributed to its role in glucose and energy metabolism ^49–52^. Our findings imply that at least some of the described TXNIP dependent effects on glucose homeostasis might also be linked to TXNIP dependent regulation of SLC7A5-SLC3A2 and the regulated import of LNAAs as cells enter and exit quiescence.

While it is clear how proliferating cells benefit from the upregulation of SLC7A5-SLC3A2, it is less clear how quiescent cells could profit from the downregulation of SLC7A5-SLC3A2. One rational for the selective endocytosis of this AA transporter could be that the cellular demand for essential AAs must be matched to lower rates of protein translation in quiescent cells. In line with this concept, also quiescent mammary epithelial cells reduced levels of V, L, I and Q ^53^. Similar results were obtained in a non-malignant murine pro-B lymphocyte cell line (FL5.12), in which the abundance of SLC3A2 at the plasma membrane was regulated by the presence of growth factors ^54^. In those cells, the consumption rate of essential AAs was decreased in G0 arrest and increased upon re-entry into the cell cycle ^55^. Hence, the downregulation of AA transporters provide means to avoid an excess/imbalance of free intracellular AAs which facilitates the controlled decrease of mTORC1 signaling and the ensuing reduction of translation rates, possibly in conjugation with the integrated stress response. Thereby, the TXNIP mediated selective downregulation of SLC7A5-SLC3A2 and the ensuing decreased import of essential AAs, helps to promote the switch from anabolic to catabolic metabolism. Moreover, the downregulation of AA transporters upon entry into quiescence might also contribute to the barrier for cellular transformation ^5^ ^56–58^.

TXNIP was specifically required for the endocytosis of SLC7A5-SLC3A2, but not for the endocytosis of SLC7A11-SLC3A2. How TXNIP interacts specifically with SLC7A5-SLC3A2 remains unclear at the moment. Yet it is clear that TXNIP requires its first PPCY motif (331) to interact with several different HECT type ubiquitin ligases, and this interaction is required for the downregulation of SLC7A5. In line with our results, SLC7A5 was shown to be ubiquitinated on N-terminal lysine residues by Nedd4-2 upon protein kinase C (PKC) activation and mTORC1 inhibition ^59,60^.

We conclude that the role of TXNIP in controlling the abundance of SLC7A5-SLC3A2 at the PM is important to match AA uptake with metabolic needs in proliferating and quiescent cells. In the future it will be important to understand how these cellular functions of TXNIP and the loss thereof translate into complex metabolic phenotypes of human patients with loss of TXNIP variants.

## Acknowledgments

We thank the patient and his family. We are grateful to Hemmo Meyer and Simona Polo for providing the YFP-tagged HECT type ubiquitin ligases. This research was funded in part by the Austrian Science Fund (FWF) (10.55776/P35874, 10.55776/P34907, 10.55776/FG20 to DT, 10.55776/P35832, 10.55776/P36600 to HF, 10.55776/P36925 to VR, 10.55776/P30196 to SH, and 10.55776/DOC82 to FH, DT, LAH, KT and MA). JK is a recipient of a DOC Fellowship of the Austrian Academy of Sciences. KT acknowledges support from the DFG (German Research Foundation, project No TH 1358/3e2), the MESI-STRAT project (grant agreement No 754688) which has received funding from the European Union’s Horizon 2020 research and innovation programme, and from the European Union European Research Council (ERC AdG BEYOND STRESS, grant agreement No 101054429) which has received funding from the European Union’s Horizon Europe research and innovation programme. For open access purposes, the author has applied a CC BY public copyright license to any author accepted manuscript version arising from this submission.

## Material and methods

### Cell culture

Patient-derived primary fibroblasts were cultured from skin biopsies. Control fibroblasts were obtained from a male, born in 2000 (WT#1) and from a male, born in 2015 (WT#2). RPE1, HEK293T, A549, H460, HCC4006, primary human fibroblasts and MEF (mouse embryonic fibroblasts) ^61^ were cultured in DMEM containing high glucose (4.5 g/L) (Sigma-Aldrich, #D6429), supplemented with 1% (vol/vol) penicillin/streptomycin (Sigma-Aldrich, #P0781) and 10% (vol/vol) FBS (fetal bovine serum) (Sigma-Aldrich, #S0615) at 37°C, in 5% CO_2_ and 98% humidity. H2228, H226, H520 and H1299 cells were cultured in RPMI 1640, glutamax supplement medium (Thermo Fisher, Gibco, # 61870044) supplemented with 1% (vol/vol) penicillin/streptomycin (Sigma-Aldrich, #P0781) and 10% (vol/vol) FBS (Sigma-Aldrich, #S0615) under the same culture conditions. Cells were passaged before reaching confluency by incubation in 1x PBS (phosphate-buffered saline, 8 g/L NaCl, 0.2 g/L KCl, 1.15 g/L Na2HPO4, 0.2 g/L KH2PO4) and 1x trypsin-EDTA solution (Sigma-Aldrich, #T4174). Cells were resuspended in growth medium and re-seeded at a 1:5 to 1:10 ratio. Cells were regularly tested negative for mycoplasma.

The AKT inhibitor MK2206 (Enzo Life Sciences, #ENZ-CHM164-0005), dissolved in DMSO (Sigma-Aldrich, #D2650), was used at a final concentration of 1 µM. The MEK inhibitor PD0325901 (Absource, #S1036), dissolved in DMSO, was used at a final concentration of 5 µM. The SLC7A5 inhibitor JPH203 (also KYT-0353, Selleckchem, #S8667), dissolved in DMSO, was used at a final concentration of 10 µM. Chloroquine (Sigma-Aldrich, #C6628) was dissolved in Millipore H2O and used at a final concentration of 12.5 µM. To inhibit dynamin mediated endocytosis, dynasore hydrate (Sigma-Aldrich, #D7693) was used as previously described ^31^ at a final concentration of 20 µM.

To maintain cells in proliferation (control cells, defined as “+ serum”), they were cultured in DMEM containing 4.5 g/L glucose and all AAs, (Sigma-Aldrich, #D6429), supplemented with 1% (vol/vol) penicillin/streptomycin (Sigma-Aldrich, #P0781) and 10% (vol/vol) FBS (Sigma-Aldrich, #S0615). To induce quiescence, cells were washed with 1x PBS and cultured in DMEM containing 4.5 g/L glucose and all AAs, supplemented with 1% penicillin/streptomycin but without FBS for 24 h (referred to as serum starvation, “-serum”).

### Proliferation assay

A similar cell number (>20000 cells) was seeded onto 6-well plates (day 0). Cells were harvested daily by trypsinization. The cell number was measured daily using a CASY Cell Counter and Analyzer (OMNI Life Science) for 4 days. The cell proliferation index was calculated as described before ^62^. The fold increase was calculated by normalizing to the cell number obtained 24 h after seeding (day 1).

For measuring cell proliferation in an automated manner, 50000 cells were seeded onto a 6-well plate and cultured at standard cell culture conditions. Cells were set on serum free medium 24 h after seeding and refed after 48 h of serum starvation. Cell proliferation was tracked using a Sartorius IncuCyte® S3 Live Cell Imaging System and confluency measurement was performed every 4 h for 5 days. Data were acquired and analyzed using the Sartorius IncuCyte® S3 software version 2022B.

### Preparation of whole cell protein extracts, SDS-PAGE and Western Blot analysis

Cells were washed in 1x cold PBS, harvested using a cell scraper and pelleted at 13.000 rpm for 5 min at 4°C. The cell pellets were resuspended in cold RIPA lysis buffer (50 mM Tris-HCl pH 7.5, 150 mM NaCl, 1% NP-40, 2 mM EDTA pH 8, 0.1% SDS, 0.5% sodium deoxycholate) or lysis buffer (50 mM Tris-HCl pH 7.5, 150 mM NaCl, 1% Triton X-100, 10% glycerol, 0.5 mM EDTA), supplemented with 50 mM NaF, 10 µg/mL leupeptin, 1 mM pefablock, 1 µg/mL pepstatin, 10 µg/mL aprotinin, 2 mM Na_3_VO_4._ Cells were lysed for 30 min on ice, centrifuged at 13.000 rpm for 10 min at 4°C and the cleared lysate was obtained. Protein concentration was measured using Micro BCA Protein Assay Kit (Thermo Scientific, #23235) according to the manufacturer’s instructions. 20 µg protein was supplemented with 5x Urea sample buffer (8 M Urea, 5% SDS, 40 mM Tris pH 6.8, 0.1 mM EDTA, 0.4 mg/mL bromphenol blue, 1% final ϕ3-mercaptoethanol) at 1x final concentration. Lysates were separated by SDS-PAGE. Proteins were transferred onto polyvinylidene fluoride (PVDF) membranes (Amersham Hybond PVDF, #10600023 – 0.45 µm) using a wet transfer system (Bio-Rad) at constant 80V for 2 h (1x transfer buffer: 20% methanol, 25 mM Tris-HCl, 192 mM glycine). Membranes were blocked in 5% milk in Tris-buffered saline buffer (TBS, 0.5 M NaCl, 20 mM Tris, pH 7.5) with 0.05% Tween 20 (TBS-T) and probed with the respective antibodies over night at 4°C. The membranes were washed with TBS-T and probed with the respective horseradish peroxidase-linked secondary antibodies (1:5000 dilution) for 1 h at RT. After washing with TBS-T, the membrane was incubated with WesternBright Chemiluminescence substrate solution (Advansta) (Biozym, #541005X). The chemiluminescence signal was detected either on X-ray films or with a CCD-Camera (Fusion FX, Vilber Lourmat). The signal was quantified with ImageJ and normalized to GAPDH. To quantify the phosphorylation of proteins, the signals were normalized to the unphosphorylated total protein levels.

### Denaturing SLC7A5-ALFA immunoprecipitation

Immunoprecipitation of SLC7A5-ALFA under denaturing lysis conditions was adopted from ^63^. Cells expressing SLC7A5-ALFA were either grown in growth medium (+ serum) or serum starved for 7 or 8 h (-serum). Cells were washed with 1x PBS and harvested. Cell pellets were resuspended in lysis buffer I (50 mM Tris pH 8, 150 mM NaCl, 50 mM NaF, 5 mM EDTA, 1% SDS, 10 nM NEM, 10 µg/mL leupeptin, 1 mM pefablock, 1 µg/mL pepstatin, 10 µg/mL aprotinin) and incubated 30 min at RT. Protein concentration was measured using Micro BCA Protein Assay Kit (Thermo Scientific, #23235) according to the manufacturer’s instructions. Equal amounts of proteins were diluted by addition of 800 µl lysis buffer II (50 mM Tris pH 8, 150 mM NaCl, 50 mM NaF, 5 mM EDTA, 1% Triton X-100, 10 nM NEM, 10 µg/mL leupeptin, 1 mM pefablock, 1 µg/mL pepstatin, 10 µg/mL aprotinin). Samples were subjected to immunoprecipitation. Beads (ALFA Selector ST, NanoTag Biotechnologies, #N1516) were equilibrate in washing buffer I (50 mM Tris pH 8, 150 mM NaCl, 1% Triton X-100). Cell suspension and beads were incubated rotating at 4°C for 3 h. Beads were recovered with a magnetic rack and washed twice with washing buffer I and three times with washing buffer II (50 mM Tris pH 8, 300 mM NaCl, 1% Triton X-100). Proteins were eluted with 5x SDS sample buffer (250 mM Tris-HCl pH 6.8, 10% SDS, 50% glycerol, 10% ϕ3-mercaptoethanol) and denatured for 10 min at 95°C. For detection of ubiquitinated proteins, samples were separated by SDS-PAGE. Proteins were transferred onto PVDF membranes (Amersham Hybond PVDF, #10600023 – 0.45 µM (300mm)) using a wet transfer system (Bio-Rad) at constant 80V for 2 h. After transfer, the membrane was blocked with 10% BSA in 0.45% Tween 20 for at least 1 h and incubated with anti-ubiquitin antibody (Santa Cruz, ##3936S) overnight in 10% BSA blocking solution.

### Non-denaturing ALFA-TXNIP immunoprecipitation

Non-denaturing immunoprecipitation of ALFA-TXNIP was adopted from ^27^. Briefly, cells were washed with 1x PBS and harvested by scraping. Cells were lysed in lysis buffer (30 mM Tris pH 7.5, 120 mM NaCl, 20 mM NaF, 0.5% octyl-ϕ3-D-glucopyranosid) (Sigma-Aldrich, #O8001) and protease inhibitors (10 µg/mL leupeptin, 1 mM pefablock, 1 µg/mL pepstatin, 10 µg/mL aprotinin, 2 mM Na_3_VO_4_). Cells were lysed for 30 min on ice, centrifuged at 13.000 rpm, at 4°C for 10 min and cleared lysate was obtained. Protein concentration was measured using Micro BCA Protein Assay Kit (Thermo Scientific, #23235) according to the manufacturer’s instructions. Equal amounts of proteins were subjected to immunoprecipitation. Beads (ALFA Selector ST, NanoTag Biotechnologies, #N1516) were equilibrated in lysis buffer. Cell suspension and beads were incubated rotating at 4°C for 3 h. After incubation beads were washed three times for 10 minutes at 4 °C using wash buffer (30 mM Tris pH 7.5, 150 mM NaCl, 0.1% octyl-ϕ3-D-glucopyranosid). Proteins were eluted in 100µl 2x Urea sample buffer at 42°C for 30 min.

### Genetic modifications and cloning

Genetic modifications were performed by PCR using standard techniques or by using the Gateway cloning method (Thermo Fisher Scientific) according to manufacturer’s instructions. All plasmids and primers used in this study are listed in Supplementary Table 3 and Supplementary Table 4. The sequence-verified plasmids were then used for lentiviral transduction of RPE1 cells. To generate virus particles, HEK293T cells were seeded in a 6-well tissue culture plate and transfected with 2 µg lentiviral vector containing the gene of interest together with virus packing vectors (1 µg pVSV-G and 1 µg psPAX2) using polyethylenimine (PEI, Polyscience, #23966-100) as transfection reagent. 4 -8 h after transfection, the medium was changed. Virus particle-containing medium was collected from the HEK293T cells 48 h and 72 h after transfection and was used to infect RPE1 cells. Polybrene was added to the virus-particle-containing medium to help lentivirus integration (1:500 dilution, Sigma-Aldrich, #107689). After 10 days of selection with 1 µg/mL blasticidin (InvivoGen, #ant-bl-5b), the efficacy of the virus transduction was tested via SDS-PAGE and Western Blot analysis.

Genetic deletion using CRISPR/Cas9 mediated gene editing was performed as previously described ^64^. For CRISPR/Cas9-mediated depletion, guide RNA (gRNA) targeting primers for SLC7A5 (5’-caccgCGTGAACTGCTACAGCGTGA -3’ and 5’-aaac TCACGCTGTAGCAGTTCACGc -3’) and TXNIP (5’-caccgGTAAGTGTGGCGGGCCACAA-3’ and 5’-aaacTTGTGGCCCGCCACACTTACc-3’). RPE1 cells were transiently transfected with the gRNA-containing pSpCas9(BB)-2A-GFP plasmid (Addgene, #48138) using Lipofectamin LTX reagent (Thermo Scientific, #15338-100) according to the manufacturer’s protocol. 24 h after transfection, GFP-positive, transfected cells were enriched by flow cytometry (BD FACSAria III Cell Sorter, BD Biosciences). The sorted cell suspension was used for the generation of single cell clones. Depletion efficiency was verified by PCR using specific primers and Western Blot analysis.

For the transient transfection of HEK293T Lipofectamine LTX & Plus reagent (Thermo Scientifc, #15338100) was used according to the manufacturer’s protocol. HEK293T cells were seeded in a 10 cm dish. At around 80% confluency, 16 µg of the respective plasmids were diluted in 500 µl of OptiMEM (Thermo Scientific, #31985062) in the presence of 48 µl of LTX-and 16 µl of Plus-reagent for the transfection. The transfected cells were incubated for 4 h before the transfection medium was discarded and replaced with 10 ml of fresh DMEM. After that, cells were incubated for 24 h under standard conditions.

### Immunofluorescence microscopy

Cells were grown on 12 mm glass coverslips (Hartenstein GmbH, #DKR1) and fixed with 4% paraformaldehyde (PFA) for 10 -15 min at RT. Immunofluorescence staining of SLC3A2 and ALFA-tagged proteins was performed as described previously ^65^. Briefly, cells were permeabilized and blocked in blocking buffer (10% FBS, 0.3% saponin (BioChemika, #84510), sodium azide (0.02% final) in 1x PBS) for 30 min at 37°C. Cells were incubated with mouse anti-SLC3A2 (BD Pharmingen, #556074) or rabbit anti-ALFA (NanoTag Biotechnologies, #N1581) in a 1:100 dilution in blocking buffer overnight at 4°C. For double-staining with LAMP1, cells were simultaneously incubated with rabbit anti-LAMP1 (Cell Signaling, #9091) in a 1:800 dilution in blocking buffer. Cells were washed once with wash solution (0.03% saponin, sodium azide (0.02% final) in 1x PBS), following three washing steps with PBS and incubated with Alexa Fluor 488 goat anti-mouse IgG (Invitrogen, #A11001) and Alexa Fluor 568 goat anti-rabbit IgG (Invitrogen, #A11011) in a 1:500 dilution in blocking buffer for 1 h at RT.

For immunofluorescence labeling of SLC1A5, cells were permeabilized in in 0.2% Triton X-100 (Thermo Scientific, #28314) in PBS for 5 min and blocked in blocking buffer (2% gelatin, 150 mM NaCl, 10 mM Pipes pH 6.8, 5 mM EGTA, 5 mM glucose, 5 mM MgCl_2_, 50 mM NH_4_Cl) for 1 h at RT. Cells were incubated with rabbit anti-SLC1A5 (Sigma-Aldrich, #HPA035240) in a 1:50 dilution in blocking buffer for 1 h at RT. For double-staining with LAMP1, cells were simultaneously incubated with mouse anti-LAMP1 (DSHB, #H4A3-S) in a 1:50 dilution in blocking buffer. Cells were washed with PBS and incubated with Alexa Fluor 488 goat anti-rabbit IgG (Invitrogen, #A11008) and Alexa Fluor 568 goat anti-mouse IgG (Invitrogen, #A11031) in a 1:500 dilution in blocking buffer for 1 h at RT.

For immunofluorescence labeling of SLC2A1, cells were permeabilized and blocked in blocking buffer (2% gelatin, 150 mM NaCl, 10 mM Pipes pH 6.8, 5 mM EGTA, 5 mM glucose, 5 mM MgCl_2_, 50 mM NH_4_Cl) containing 0.025% saponin (BioChemika, #84510) for 1 h at RT. Cells were incubated with rabbit anti-SLC2A1 (Merck, #07-1401) in a 1:200 dilution in blocking buffer for 1 h at RT. Cells were washed with PBS and incubated with Alexa Fluor 488 goat anti-rabbit IgG (Invitrogen, #A11008) in a 1:500 dilution in blocking buffer for 1 h at RT. Cells were washed with 1x PBS, incubated with DAPI (Sigma-Aldrich, #D9542) in a 1:20000 dilution in 1x PBS and mounted in Mowiol (Sigma-Aldrich, #81381).

Confocal microscopy was performed on a LSM980 AiryScan 2 (ZEISS). Confocal Z-series stack (step size 0.22 µm) images were acquired using the AiryScan detector, a LD LCI Plan-Apochromat 63x / 1.2 with glycerol immersion objective (ZEISS) and the 405 nm, 488 nm and 561 nm laser lines. To compare signals of a protein of interest in proliferating, quiescent or treated cells, laser settings were kept constant while imaging both conditions. The ZEISS Zen 3.5 software was used as recording software and for processing of the raw images (AiryScan processing). Brightness and contrast were adjusted using ImageJ software. For merged images, the levels of red, green and blue channels were separately adjusted.

### Cell surface staining of SLC3A2 and FACS analysis

To measure the difference in surface staining of SLC3A2 between proliferating and quiescent cells, cells were either grown in growth medium or serum starved for 24 h before the experiment. The staining was performed as previously described ^65^. Cells were washed with PBS, trypsinized and resuspended in FACS buffer (10% FBS in 1x PBS) containing anti-SLC3A2 antibody (BD Pharmingen, #556074) at a dilution of 1:100. The antibody was incubated for 30 min on ice. Cells were centrifuged at 1000 rpm for 5 min at 4°C and resuspended in FACS buffer containing Alexa Fluor 488 goat anti-mouse IgG (Invitrogen, #A11001) at a dilution of 1:500. The antibody was incubated for 30 min on ice in the dark. Cells were centrifuged at 1000 rpm for 5 min at 4°C and resuspended in 200µl FACS buffer. The fluorescence intensity was measured by flow cytometry (Attune NxT Flow Cytometer, Life Technologies). For every experiment, more than 3500 cells were analyzed. Flow cytometry analysis was performed with FlowJo software 10.8.1 (BD Life Sciences). Cells were gated by forward versus side scatter (FSC-A vs. SSC-A) to identify cell population, forward scatter width versus forward scatter area (FSC-W vs FSC-A) for doublet exclusion and fluorescence intensity (BL1-A) vs cell count to reflect Alexa-488 signal populations. Unstained cells were excluded from analysis.

### DNA content for cell cycle analysis of fixed cells stained with propidium iodide (PI)

Cells were washed with PBS and harvested by trypsinization. Cells were centrifuged at 1000 rpm for 5 min at 4°C. The pellet was washed in cold PBS. Cells were fixed by resuspending the cells in 1 ml ice cold 70% vol/vol ethanol in PBS. Samples were stored at -20°C for at least 30 min. The ethanol was washed away with ice cold PBS twice. Cells were then incubated with propidium iodide (PI) at a final dilution of 1:25 and RNAse A (Thermo Scientific, #EN0531) at a final dilution of 1:100 in PBS, for 30 min at 37°C. The fluorescence intensity was measured by flow cytometry (Attune NxT Flow Cytometer, Life Technologies). For every experiment, more than 1500 cells were analyzed. Data were analyzed using ModFit LT (Verity Software House). Cells were gated by forward versus side scatter (FSC-A vs. SSC-A) to identify cell population. PI-positive cells were gated for doublet exclusion by width vs height (BL3-H vs BL3-W) and fluorescence intensity (BL3-A) vs cell count to reflect PI-positive signal populations.

RNA was extracted from cells using the RNeasy Mini Kit (Quiagen, #74104) according to the manufacturer’s instructions. 800 ng RNA were reverse transcribed into cDNA using the LunaScript RT Super Mix Kit (NEB, #E3010) according to manufacturer’s protocol. For mRNA fold regulation analysis, quantitative real-time PCR was performed using gene specific primers listed in Supplementary Table 5. The Biozym Blue S’Green qPCR Kit (Biozym, #F-415S) was used according to manufacturer’s protocol. All primers were synthesized by Sigma-Aldrich and diluted in Millipore H_2_O to a concentration of 10 µM. Each condition was performed in technical triplicates from three biological replicates. Fold regulation was calculated using the 2-^1¢1¢Ct^ method and normalized to the housekeeping gene *ribosomal protein lateral stalk subunit P0 (RPLP0)*.

### Uptake of radioactive labeled AAs

To analyze the uptake of L-glutamine, cells were washed with PBS and incubated with 0.3 µCi/mL [^14^C_5_]-L-glutamine (Hartmann Analytic, #MC1124) in glutamine-free medium (Sigma-Aldrich, #D6546). To analyze the uptake of L-leucine, cells were washed with PBS and incubated with 0.5 µCi/mL [4,5-^3^H]-L-Leucine (Hartmann Analytic, #MT672E) in AA free medium (USBiological, #D9811-01). The uptake of L-isoleucine was initiated by incubating the cells with 0.005 µCi/mL [^14^C_5_]-L-isoleucine (Hartmann Analytic; #MC174) in AA free medium. The uptake was performed for 15 min at 37°C. To stop the reaction, cells were washed twice with ice cold PBS. Cells were harvested by trypsinization and the cell pellet was lysed in 100 µl lysis buffer (50 mM Tris-HCl pH 7.5, 150 mM NaCl, 1% NP-40, 2 mM EDTA pH 8, 0.1% SDS, 0.5% sodium deoxycholate) for 30 min on ice. Cells debris were removed by centrifugation (13000 rpm, 4°C, 10 min) and the supernatant was analyzed by liquid scintillation counting (LSC-Universal cocktail, Roth, #0016.3; HITACHI AccuFlex LSC-800). AA uptake was calculated as counts per min (cpm) normalized to the protein concentration (µg) of each cleared lysate. Protein concentration was determined by using Micro BCA Protein Assay Kit (Thermo Scientific, #23235) according to the manufacturer’s instructions.

### Glucose uptake assay

The glucose uptake assay was performed as described previously ^66^. Briefly, glucose uptake was initiated by adding 2-NBDG (2-(N-(7-Nitrobenz-2-oxa-1,3-diazol-4-yl)amino)-2-deoxyglucose) (AAT Bioquest, #36702, dissolved in DMSO) at a final concentration of 100 µM in the respective medium. To inhibit glucose uptake, cells were simultaneously treated with 1 mM phloretin (Sigma-Aldrich, #P7912, dissolved in 50% ethanol) for 45 min. After 45 min incubation at 37°C in the dark, cells were washed with ice cold PBS to stop the reaction, trypsinized and resuspended in PBS. Cells were centrifuged at 1000 rpm for 5 min at 4°C and the pellet was resuspended in 200 µl PBS and 10% FBS. The fluorescence intensity was measured by flow cytometry (Attune NxT Flow Cytometer, Life Technologies). For every experiment, more than 5000 cells were analyzed. Flow cytometry analysis was performed with FlowJo software 10.8.1 (BD Life Sciences). Cells were gated by forward versus side scatter (FSC-A vs. SSC-A) to identify cell population, forward scatter width versus forward scatter area (FSC-W vs FSC-A) for doublet exclusion and fluorescence intensity (BL1-A) vs cell count to reflect the uptake of 2-NBDG.

### Incorporation of ^35^S labeled L-Cysteine and L-Methionine

150000 cells were seeded and after two days, these cells were left untreated or serum starved for 24 h or serum starved and then refeed for 24 h. Prior to radiolabeling the cells, the serum starved cells were washed with PBS and incubated for 30 min in L-cysteine, L-methionine, L-glutamine free medium (Sigma-Aldrich #D0422), supplemented with 2 mM L-glutamine (Gibco #25030-024) and 1% (vol/vol) penicillin/streptomycin. The proliferating cells were incubated for 30 min in L-cysteine, L-methionine, L-glutamine free medium, supplemented with 2 mM L-glutamine, 10% dialysed FBS (Thermo Fisher Scientific #26400044) and 1% (vol/vol) penicillin/streptomycin. Cells were washed with PBS, brought into suspension using 1x trypsin-EDTA solution and resuspended in the corresponding L-cysteine, L-methionine free medium. 17.6 µCi/mL [^35^S]-methionine [^35^S]-cysteine (Hartmann Analytic #21635410) was added to the cell suspension for 2 min and quenched with 10 µg/ml cycloheximide. Cell pellets were lysed for 30 min on ice, with RIPA lysis buffer, supplemented with protease inhibitors and 10 µg/ml cycloheximide. Equal amounts of protein were subjected to SDS-PAGE, (de)stained with Coomassie and dried for 1 h at 70°C. The dried gel was exposed to a phosphor-imager for 48 h. Detection of a [^35^S]-methionine [^35^S]-cysteine radiogram was performed using a Typhoon FLA 9500 detection system.

### Mass spectrometry analysis of intracellular free AAs

The analysis of intracellular AAs by mass spectrometry was performed as described previously ^67^ ^68^. Briefly, 10^6^ cells were seeded and serum starved for 24 h prior the extraction. Then, cells were washed three times with ice cold 1xPBS and extracted by adding 500 µl ice cold methanol (99,9% vol/vol, Fisher Chemicals) and 500 µl cold Millipore H_2_O, containing the internal standard (1:100), a stable isotope-labeled canonical AA mix composition (Cambridge Isotope Laboratory, #MSK-CAA-1). Cells were scraped and 500 µl ice cold chloroform (99,9% vol/vol, Sigma-Aldrich, #650498) was added to the cell extracts. Cells were agitated in a tube shaker for 20 min at 1400 rpm at 4°C, followed by centrifugation for 5 min at 16100xg at 4°C. The upper, polar phase was separated, evaporated by using a SpeedVac (Savant SPD111V SpeedVac Concentrator, Thermo Scientific) and stored at -80°C. The interphase was used to determine the protein concentration. Therefore, the interphase was evaporated and the pellet was resuspended in denaturing buffer (1.6 M Urea, 100 mM AmBiCa, pH 8.2) and sonicated for 10 sec to break up protein aggregates. Protein determination was done by using Micro BCA Protein Assay Kit (Thermo Scientific, #23235) according to the manufacturer’s instructions. For absolute quantification, AAs were analyzed by targeted LC-MS using mixed-mode chromatography (RP-AEX, reversed phase–anion exchange chromatography) and ultra-high sensitive orbitrap technology (Orbitrap Q Exactive HF-X and Orbitrap Fusion Lumos, both Thermo Scientific), operating in parallel reaction monitoring (PRM) mode. Bioinformatics data processing and statistical analysis was performed using TraceFinder (Thermo Scientific) and in-house R and Python scripts.

### Statistical analysis and software

Software used in this study, if not already specified, were Affinity Designer Version 1.10.5 (Serfi, RRID:SCR_016952), Image J Version 1.53t (RRID:SCR_003070), GraphPad Prism 9 Version 9.4.1 (458) (RRID:SCR_002798), Benchling (RRID:SCR_013955), FlowJo Version 10.8.1 (RRID:SCR_008520), ModFit LT (RRRID:SCR_016106), Primer3 Version 4.1.0 (RRID:SCR_003139), PrimerBank (RRID:SCR_006898), CHOPCHOP (RRID:SCR_015723), ZEISS ZEN Digital Imaging for Light Microscopy Version 3.5 (RRID:SCR_013672), and online tools TIDE (tracking of indels by Decomposition) (Brinkman et al. 2014), ICE Analysis online tool (Synthego, v2.0; Synthego, USA) and PrimerQuest^TM^ Tool (Integrated DNA Technologies, IDT, USA)

Results were represented as mean with standard deviation (SD) analyzed by GraphPad Prism. Statistical significance between different biological samples was determined by either paired Student’s t test or one-way/ two-way ANOVA with Tukey test for multiple comparisons, if not described otherwise.

---

**Supplementary Figure 1.**
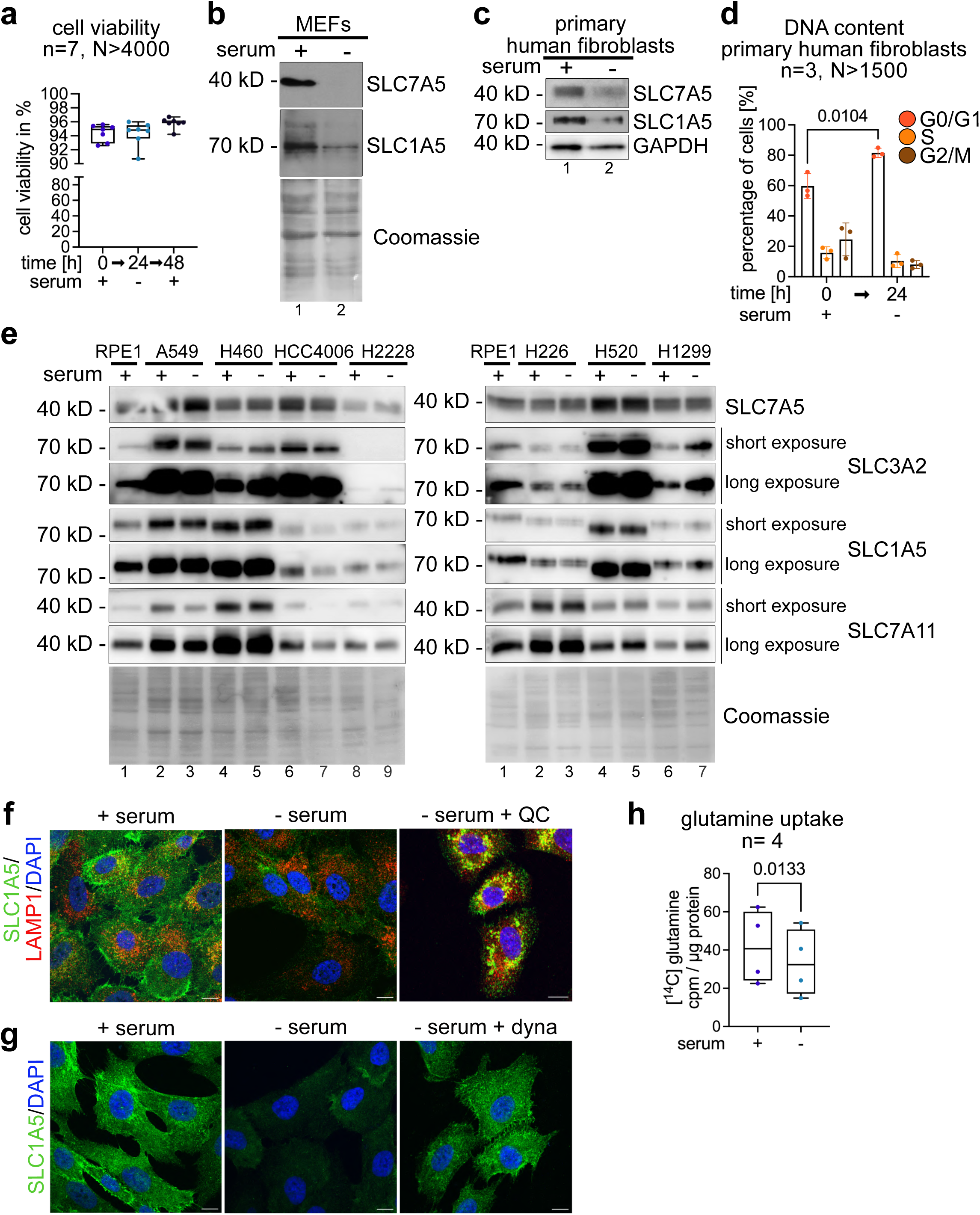
**(a)**Cell viability determined with CASY counter, before and after serum starvation and after re-addition of serum (n=7, N>4000 cells). **(b)** Immortalized MEFs were grown in growth medium with serum (+ serum) or serum starved for 24 h. Total cell lysates were analyzed by SDS-PAGE and WB with the indicated antibodies. **(c, d)** Primary human fibroblasts were grown in growth medium (+ serum) or serum starved for 24 h. **(c)** Total cell lysates were analyzed by SDS-PAGE and WB with the indicated antibodies. **(d)** Cells were harvested, permeabilized and fixed. The DNA content was analyzed by PI staining and FACS (n=3, N>1500 cells, two-way ANOVA, Tukey’s multiple comparisons test). **(e)** Total cell lysates of the indicated cell lines were analyzed by SDS-PAGE and WB with the indicated antibodies. (**f, g)** Indirect IF of PFA fixed cells stained for **(f, g)** SLC1A5 (green), **(f)** LAMP1 (red) and DAPI (blue) were analyzed by confocal microscopy. The merged images show a single plane of a Z-stack. **(f)** Incubation with 12,5 µM chloroquine (CQ) in absence of serum for 14 h (-serum, + CQ). Scale bar=10 µm **(g)** Incubation with 20 µM dynasore (dyna) in absence of serum for 14 h (-serum, + dyna). Scale bar=10 µm. **(h)** Cells were incubated with [^14^C]-glutamine for 15 min, washed and lysed. Total cell lysates were analyzed by scintillation counting. Cpm values were normalized to total protein content (cpm / µg protein, n=4, paired t-test).

**Supplementary Figure 2.**
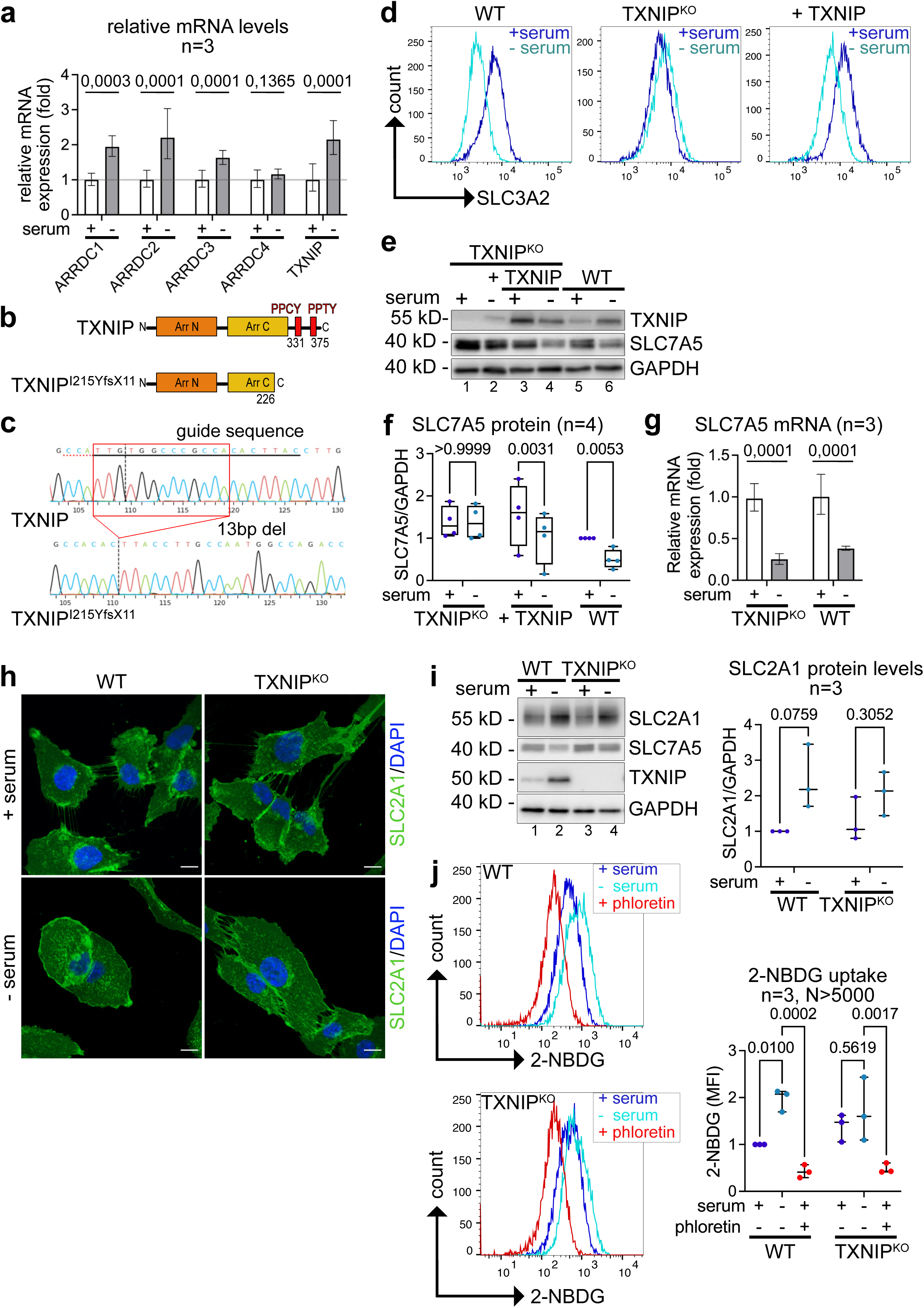
**(a, g)** qPCR analysis of **(a)** α-arrestins mRNA or **(g)** SLC7A5 mRNA in growing (+ serum) or serum starved cells (-serum). Expression levels were normalized to the housekeeping gene *RPLP0* (n=3, paired t-test). **(b)** Schematic presentation of the mutation introduced into TXNIP by gene editing. **(c)** Representative chromatogram of Sanger sequencing from the genome amplicon of WT TNIXP and gene edited TXNIP cells. The guide RNA and the protospacer adjacent motif (PAM) sequence are indicated. The vertical lane shows the Cas9 cleavage site causing a 13 base pair deletion in *TXNIP* (red box). Image was generated by the ICE Analysis online tool (Synthego, v2.0; Synthego, USA). **(d)** Representative histograms of SLC3A2 cell surface FACS from WT, TXNIP^KO^ and TXNIP^KO^ reconstituted with TXNIP cells. **(e)** WT, TXNIP^KO^ and TXNIP^KO^ reconstituted with TXNIP were grown in growth medium (+ serum) or serum starved for 24 h (-serum). Total cell lysates were analyzed by SDS-PAGE and WB with the indicated antibodies. **(f)** WB quantification of SLC7A5 protein levels, normalized to GAPDH (n=4, two-way ANOVA, Sidak’s multiple comparisons test). **(h)** Indirect IF of PFA fixed WT and TXNIP^KO^ cells stained for SLC2A1 (green) and DAPI (blue) was analyzed by confocal microscopy. The merged images show a single plane of a Z-stack. Scale bar=10 µm **(i)** Total cell lysates were analyzed by SDS-PAGE and WB with the indicated antibodies. For WB quantification, SLC2A1 protein levels were normalized to GAPDH (n=3, two-way ANOVA, Sidak’s multiple comparisons test). **(j)** WT and TXNIP^KO^ cells were assessed for glucose uptake by incorporation of 100 µM 2-NBDG for 45 min. Simultaneously, cells were treated with 1 mM phloretin. The fluorescence intensity was analyzed by FACS (n=3, N>5000 cells, two-way ANOVA, Sidak’s multiple comparisons test).

**Supplementary Figure 3.**
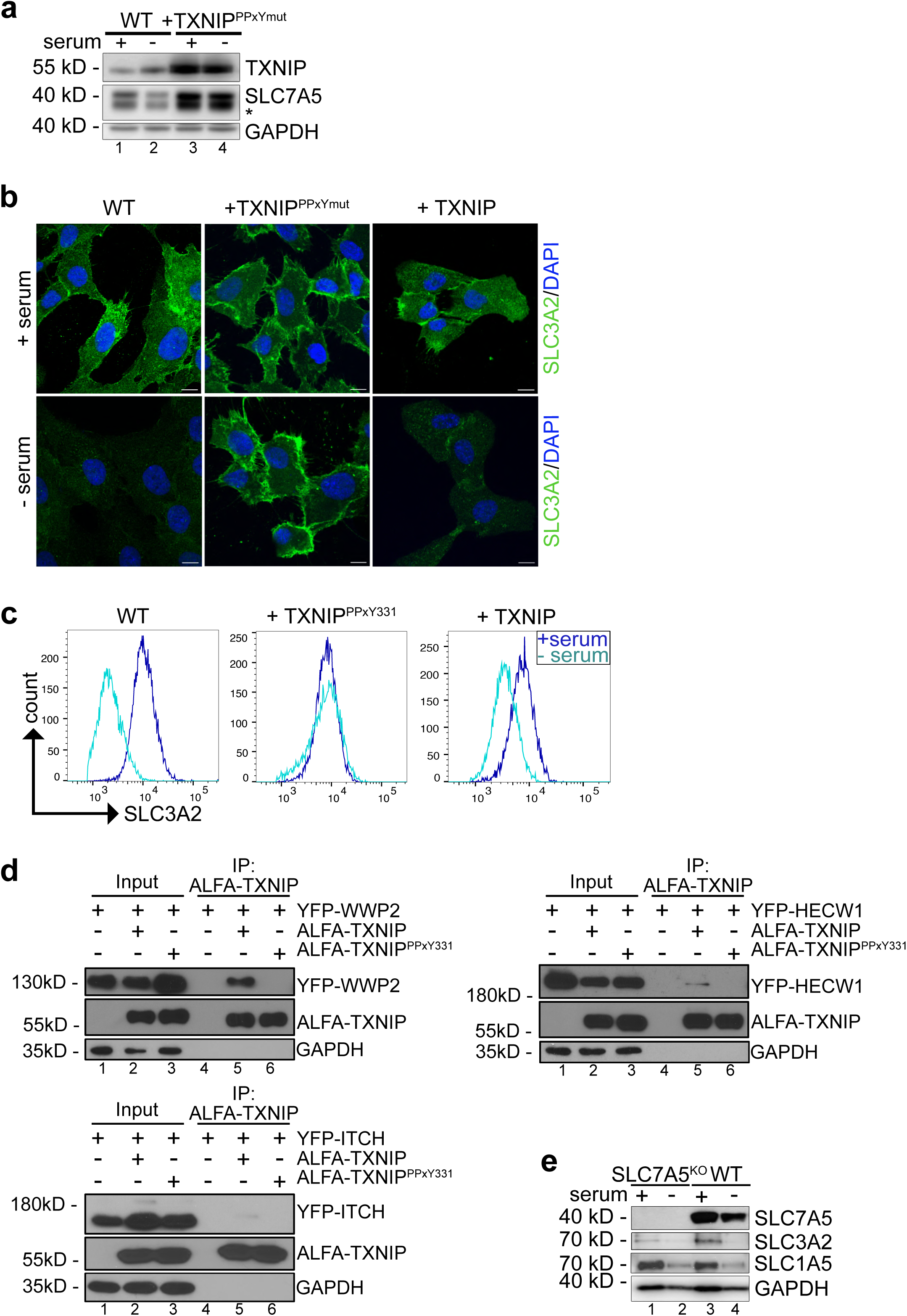
**(a)** Total cell lysates from WT or TXNIP^KO^ cells reconstituted with TXNIP^PPxYmut^ mutants were analyzed by SDS-PAGE and WB with the indicated antibodies. The asterisk (*) labels a second band that was only detected using the monoclonal SLC7A5 antibody. **(b)** Indirect IF of PFA fixed cells (WT, TXNIP^KO^ cells reconstituted with TXNIP or TXNIP^PPxYmut^) stained for SLC3A2 (green) and DAPI (blue) was analyzed by confocal microscopy. The images show a single plane of a Z-stack. Scale bar = 10 µm. **(c)** Representative histograms of SLC3A2 cells surface FACS of the indicated cells. **(d)** SDS-PAGE and WB analysis with the indicated antibodies from non-denaturing ALFA-immunoprecipitations (IP) from HEK293T cells expressing ALFA-TXNIP or the mutated ALFA-TXNIP^PPxY331^ variant and the indicated YFP-tagged HECT-type ubiquitin ligase. **(e)** SDS-PAGE and WB with the indicated antibodies of SLC7A5^KO^ and WT cells.

**Supplementary Figure 4.**
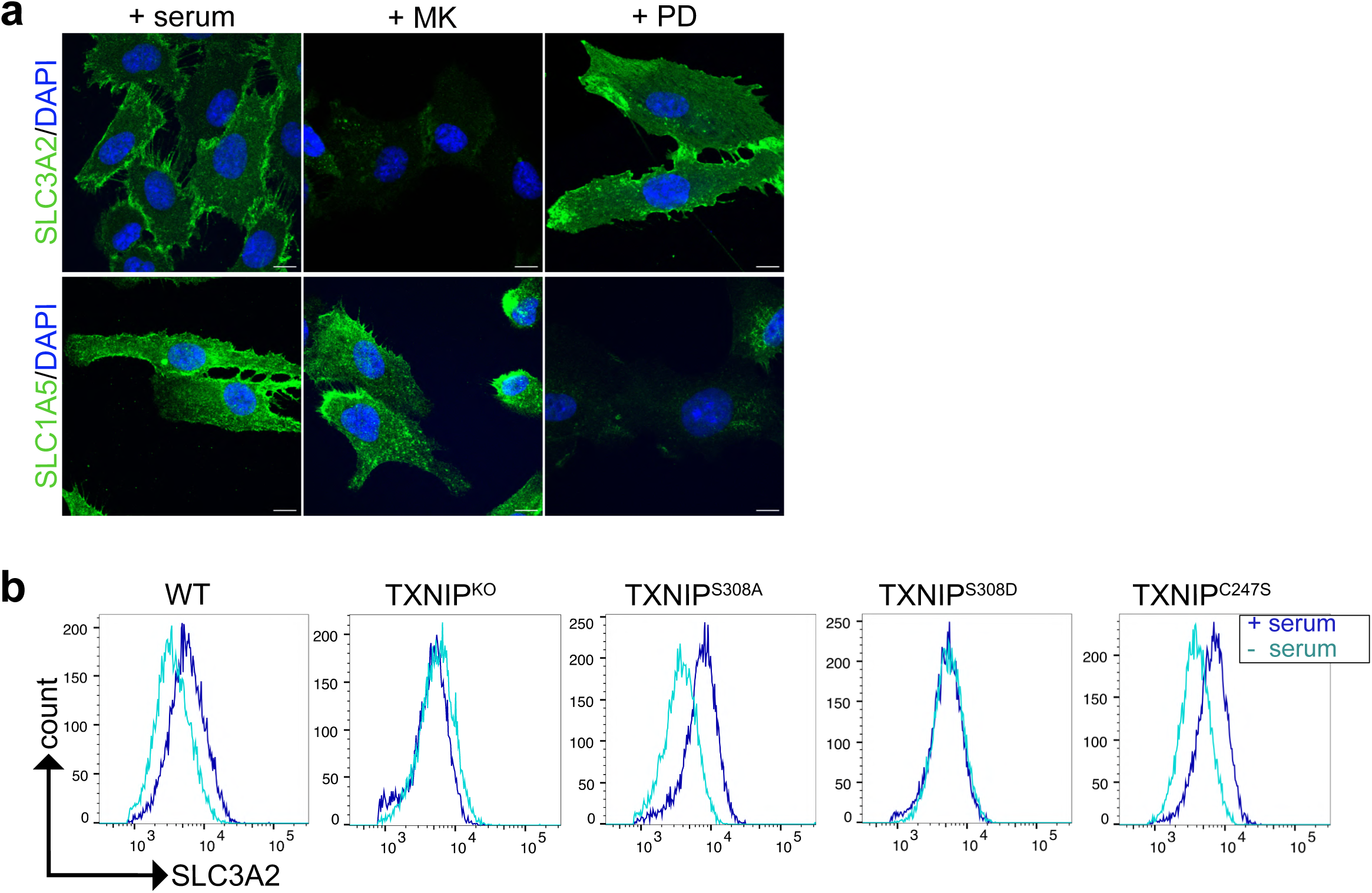
**(a)** Indirect IF of PFA fixed cells before or after serum starvation or incubation with 1 µM MK2206 (MK) or 5 µM PD0325901 (PD) stained for SLC3A2 (green, upper row) or SLC1A5 (green, lower row), and DAPI (blue). Cells were analyzed by confocal microscopy. The images show a single plane of a Z-stack. Scale bar = 10 µm. **(b)** Representative histograms of SLC3A2 cells surface FACS of the indicated cells.

**Supplementary Figure 5.**
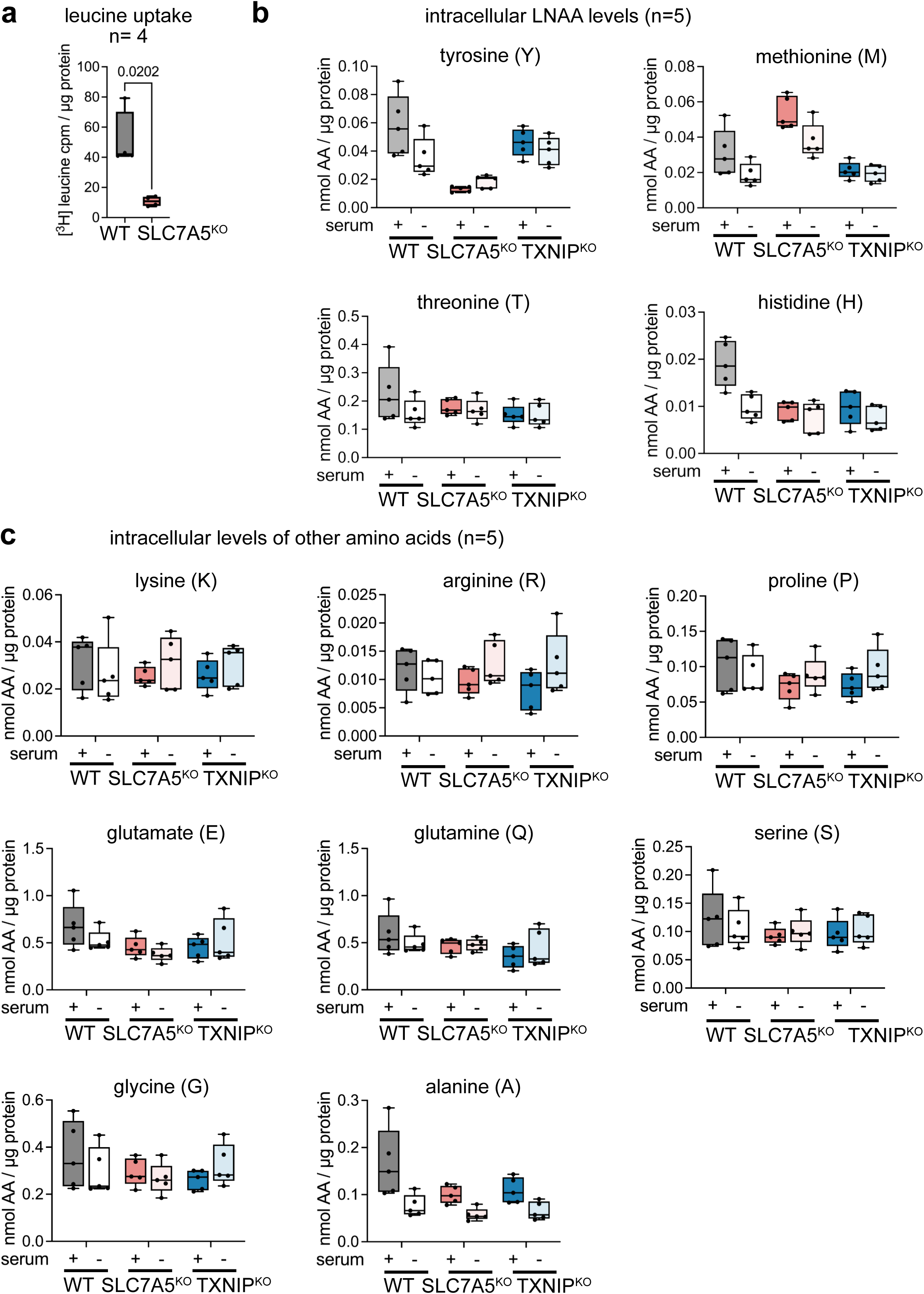
**(a)** WT and SLC7A5^KO^ cells were incubated with [^3^H]-leucine for 15 min, washed and lysed. Cell lysates were analyzed by scintillation counting. Cpm values were normalized to total protein content (n=4, paired t-test). **(b, c)** WT, SLC7A5^KO^ and TXNIP^KO^ cells were grown in growth medium (+ serum) or serum starved for 24 h (-serum). Mass spectrometry analysis of free AAs, normalized to total protein content (nmol AA / µg protein, n=5).

**Supplementary Table 1.** AA, glucose and lactate levels in blood plasma of the TXNIP deficient patient over the indicated years.

| **AA [μmol/l]** | **06/2014** | **07/2014** | **Reference values** | **02/2016** | **03/2018** | **11/2021** | **09/2022** | **Reference values** |
| --- | --- | --- | --- | --- | --- | --- | --- | --- |
| leucine (L) | 60.51 | 175.52 | 46 - 109 | 346.58 | 226.89 | 212.26 | 141.50 | 45 - 155 |
| Isoleucine (I) | 13.38 | 91.56 | 26 - 53 | 173.06 | 132.25 | 129.59 | 82.64 | 26 - 94 |
| tryptophan (W) | 25.21 | 51.44 | n.a | 76.35 | 45.51 | 40.21 | 41.48 | n.a |
| valine (V) | 99.97 | 148.00 | 57 - 262 | 529.46 | 376.67 | 389.20 | 229.86 | 57 - 270 |
| phenylalanine (F) | 46.45 | 41.50 | 23 - 110 | 84.41 | 54.64 | 53.02 | 56.75 | 23 - 70 |
| methionine (M) | 12.14 | 24.43 | 3 - 41 | 36.19 | 18.77 | 10.83 | 12.30 | 3 - 29 |
| proline (P) | 191.78 | 197.21 | 52 - 277 | 264.22 | 228.78 | 214.37 | 275.47 | 51 - 185 |
| tyrosine (Y) | 38.54 | 52.02 | 11 - 112 | 110.43 | 70.10 | 51.76 | 50.47 | 11 - 120 |
| Fischer's ratio (FR) | 2.05 | 4.46 | 2.10 - 4 | 5.38 | 5.90 | 6.98 | 4.23 | 2.10 - 4 |
| **glucose [mg/dl]** | 34 | 71 | > 50 | 88 * | 78 | 86 | 103 | > 50 |
| **lactate [mg/dl]** | 68.30 | 46 | 5 - 22 | 24.33 * | 50.5 | 26 | 56 | 5 - 22 |

\* The glucose and lactate values were collected on a different day than the AA levels.

**Supplementary Table 2.** Summary of antibodies, reagents and software used in this study.

| Reagents/resource | Reference or source | Identifier or catalog number | Additional information |
| --- | --- | --- | --- |
| **Antibodies** |  |  |  |
| SLC1A5/ASCT2 | Merck | #abn73,<br>RRID:AB_10807715 | WB (1:1000) |
| SLC1A5/ASCT2 (V501) | Cell Signaling | #5345,<br>RRID:AB_10621427 | WB (1:1000) |
| SLC1A5/ASCT2 | Sigma-Aldrich | #HPA035240,<br>RRID:AB_10604092 | IF (1:50) |
| SLC7A5/LAT1 (polyclonal) | Cell Signaling | #5347S,<br>RRID:AB_10695104 | WB (1:2000) |
| SLC7A5/LAT1 (E9O4D) (monoclonal) | Cell Signaling | #32683 | WB (1:2000) |
| SLC7A5/LAT1 (D10) | Santa Cruz | #sc-374232,<br>RRID:AB_10988206 | WB (1:100) |
| SLC3A2/4F2hc (D6O3P) | Cell Signaling | #13180S,<br>RRID:AB_2687475 | WB (1:1000) |
| SLC3A2/4F2hc | BD Biosciences | #556074,<br>RRID:AB_396341 | IF, FACS (1:100) |
| SLC2A1/Glut1 | Merck | #07-1401,<br>RRID:AB_1587074 | WB (1:2000), IF (1:200) |
| SLC38A2/SNAT2 | MBL International | #BMP081,<br>RRID:AB_10597880 | WB (1:100) |
| SLC7A11/xCT (D2M7A) | Cell Signaling | #12691,<br>RRID:AB_2687474 | WB (1:1000) |
| TXNIP (D5F3E) | Cell Signaling | #14715S,<br>RRID:AB_2714178 | WB (1:2000) |
| p-S6K<sup>T389</sup> (108D2) | Cell Signaling | #9234S,<br>RRID:AB_2269803 | WB (1:1000) |
| S6K | Cell Signaling | #9202S,<br>RRID:AB_331676 | WB (1:1000) |
| p-S6<sup>S240/244</sup> | Cell Signaling | #2215S,<br>RRID:AB_331682 | WB (1:2000) |
| S6 (54D2) | Cell Signaling | #2317S,<br>RRID:AB_2238583 | WB (1:1000) |
| p-ERK1/2<sup>T202/Y204</sup> (D13.14.4E) | Cell Signaling | #4370S,<br>RRID:AB_2315112 | WB (1:1000) |
| ERK1/2 | Cell Signaling | #9102S,<br>RRID:AB_330744 | WB (1:1000) |
| p27<sup>Kip1</sup> (D69C12) | Cell Signaling | #3686S,<br>RRID:AB_2077850 | WB (1:1000) |
| p-AKT<sup>S473</sup> (D9E) | Cell Signaling | #4060S,<br>RRID:AB_2315049 | WB (1:1000) |
| p-AKT<sup>T308</sup> | Cell Signaling | #9275,<br>RRID:AB_329828 | WB (1:1000) |
| AKT | Cell Signaling | #9272S,<br>RRID:AB_329827 | WB (1:1000) |
| ALFA | NanoTag Biotechnologies | #N1581, RRID:AB_3075997 | WB (1:1000), IF (1:100) |
| Transferrin Receptor (Tfr) | DSHB | #G1/221/12, RRID:AB_2201506 | WB (1:1000) |
| EGF Receptor (EGFR) (1005) | Santa Cruz | #sc-03-G | WB (1:500) |
| GAPDH (D4C6R) | Cell Signaling | #97166S, RRID:AB_2756824 | WB (1:5000) |
| LAMP1 | DSHB | #H4A3, RRID:AB_2296838 | IF (1:50) |
| LAMP1 (D2D11) | Cell Signaling | #9091, RRID:AB_2687579 | IF (1:800) |
| NEDD4 | Cell Signaling | #5344S, RRID:AB_10560514 | WB (1:1000) |
| WWP1 | Cell Signaling | #70140S, RRID:AB_3662810 | WB (1:1000) |
| WWP2 | Cell Signaling | #41182S, RRID:AB_3662809 | WB (1:1000) |
| ITCH | Cell Signaling | #12117S, RRID:AB_2797822 | WB (1:1000) |
| GFP | Roche Life Science | (Roche Cat# 11814460001, RRID:AB_390913) | WB (1:1000) |
| Ubiquitin (P4D1) | Santa Cruz | #3936S, RRID:AB_628423 | WB (1:500) |
| Goat anti-mouse IgG-peroxidase | Sigma-Aldrich | #A4416, RRID:AB_258167 | WB (1:5000) |
| Goat anti-rabbit IgG-peroxidase | Sigma-Aldrich | #A0545, RRID:AB_257896 | WB (1:5000) |
| Alexa Fluor 488 goat anti-rabbit IgG | Invitrogen | #A11008, RRID:AB_143165 | IF (1:500) |
| Alexa Fluor 488 goat anti-mouse IgG | Invitrogen | #A11001, RRID:AB_2534069 | IF (1:500) |
| Alexa Fluor 568 goat anti-mouse IgG | Invitrogen | #A11031, RRID:AB_144696 | IF (1:500) |
| Alexa Fluor 568 goat anti-rabbit IgG | Invitrogen | #A11011, RRID:AB_143157 | IF (1:500) |
| **Chemicals** |  |  |  |
| Blasticidin | InvivoGen | #ant-bl-5b |  |
| High glucose Dulbecco's modified Eagle's medium (DMEM) | Sigma-Aldrich | #D6429 |  |
| FSB | Sigma-Aldrich | #S0615 |  |
| Trypsin-EDTA solution | Sigma-Aldrich | #T4174 |  |
| MK2206 | Enzo Life Sciences | #ENZ-CHM164-0005 |  |
| PD0325901 | Absource | #S1036 |  |
| JPH203 | Selleckchem, | #S8667 |  |
| chloroquine | Sigma-Aldrich | #C6628 |  |
| dynasore hydrate | Sigma-Aldrich | #D7693 |  |
| penicillin/streptomycin | Sigma-Aldrich | #P0781 |  |
| WesternBright<br>Chemiluminescence<br>substrate solution<br>(Advansta) | Biozym | #541005X |  |
| Lipofectamin LTX<br>reagent | Thermo Scientific | #15338-100 |  |
| polyethylenimine (PEI) | Polyscience | #23966-100 |  |
| Polybrene | Sigma-Aldrich | #107689 |  |
| DAPI | Sigma-Aldrich | #D9542 |  |
| Mowiol | Sigma-Aldrich | #81381 |  |
| saponin | BioChemika | #84510 |  |
| RNAse A | Thermo Scientific | #EN0531 |  |
| [<sup>14</sup>C<sub>5</sub>]-L-glutamine | Hartmann Analytic | #MC1124 |  |
| [4,5-<sup>3</sup>H]-L-Leucine | Hartmann Analytic | #MT672E |  |
| [<sup>14</sup>C<sub>5</sub>]-L-Isoleucine | Hartmann Analytic | #MC174 |  |
| glutamine-free DMEM | Sigma-Aldrich | #D6546 |  |
| AA free DMEM | USBiological | #D9811-01 |  |
| 2-NBDG | AAT Bioquest | #36702 |  |
| phloretin | Sigma-Aldrich | #P7912 |  |
| stable isotope-labeled<br>canonical AA mix<br>composition | Cambridge Isotope<br>Laboratory | #MSK-CAA-1 |  |
| Cycloheximide | Sigma | Cat. # C7698 |  |
| anti-ALFA selector ST<br>magnetic beads | NanoTag<br>Biotechnologies | Cat. # N1516 |  |
| **Critical commercial**<br>**assays** |  |  |  |
| Micro BCA Protein<br>Assay Kit | Thermo Scientific | #23235 |  |
| RNeasy Mini Kit | Quiagen | #74104 |  |
| LunaScript RT Super<br>Mix Kit | NEB | #E3010 |  |
| Biozym Blue S'Green<br>qPCR Kit | Biozym | #F-415S |  |
| **Software, algorithm** |  |  |  |
| Image J |  | Version 1.53t,<br>RRID:SCR_003070 |  |
| Affinity Designer | Serfi | Version 1.10.5,<br>RRID:SCR_016952 |  |
| GraphPad Prism 9 |  | Version 9.4.1,<br>RRID:SCR_002798 |  |
| Benchling |  | RRID:SCR_013955 |  |
| Flow Jo |  | Version 10.8.1,<br>RRID:SCR_008520 |  |
| ModFit LT |  | RRRID:SCR_016106 |  |
| Primer3 |  | Version 4.1.0,<br>RRID:SCR_003139 |  |
| PrimerBank |  | RRID:SCR_006898 |  |
| CHOPCHOP |  | RRID:SCR_015723 |  |
| ZEISS ZEN Digital<br>Imaging for Light<br>Microscopy |  | Version 3.5,<br>RRID:SCR_013672 |  |
| PrimerQuest™ Tool | Integrated DNA<br>Technologies, IDT |  |  |
| ICE Analysis online tool | Synthego, USA | Synthego, v2.0 |  |

**Supplementary Table 3:** List of oligonucleotides used in this study.

| Name | Vector backbone | Origin |
| --- | --- | --- |
| TXNIP | pCCL-EIF1α-BlastiR-DEST | This study |
| ALFA-TXNIP | pCCL-EIF1α-BlastiR-DEST | This study |
| SLC7A5-ALFA | pCCL-EIF1α-BlastiR-DEST | This study |
| TXNIP<sup>S308A</sup>/ALFA-TXNIP<sup>S308A</sup> | pCCL-EIF1α-BlastiR-DEST | This study |
| TXNIP<sup>S308D</sup>/ALFA-TXNIP<sup>S308D</sup> | pCCL-EIF1α-BlastiR-DEST | This study |
| TXNIP<sup>C247S</sup>/ALFA-TXNIP<sup>C247S</sup> | pCCL-EIF1α-BlastiR-DEST | This study |
| ALFA-TXNIP<br>p.Ile215TyrfsTer59 | pCCL-EIF1α-BlastiR-DEST | This study |
| ALFA-TXNIP<sup>PPxY331AAxA,PPxY375AAxA</sup> | pCCL-EIF1α-BlastiR-DEST | This study |
| ALFA-TXNIP<sup>PPCY331AACA</sup> | pCCL-EIF1α-BlastiR-DEST | This study |
| pVSV-G |  | Clontech, #631530 |
| psPAX2 |  | Gift from Stephan Geley |
| pDONR-ALFA-MCS |  | Generated by Michael Widerin |
| pDONR-MCS-ALFA |  | Generated by Michael Widerin |
| pDONR-221 |  | Invitrogen, #12536017 |
| pCCL-EIF1α-BlastiR-DEST | | Gift from Stephan Geley |
| pSpCas9(BB)-2A-GFP (PX458) |  | Addgene, #48138 |
| YFP-NEDD4 | pCR3.1 | <sup>41</sup>, <sup>42</sup> |
| YFP-ITCH | pCR3.1 | <sup>41</sup>, <sup>42</sup> |
| YFP-WWP1 | pCR3.1 | <sup>41</sup>, <sup>42</sup> |
| YFP-WWP2 | pCR3.1 | <sup>41</sup>, <sup>42</sup> |
| YFP-HECW1 | pCR3.1 | <sup>41</sup>, <sup>42</sup> |
| YFP-HECW2 | pCR3.1 | <sup>41</sup>, <sup>42</sup> |

**Supplementary Table 4:**
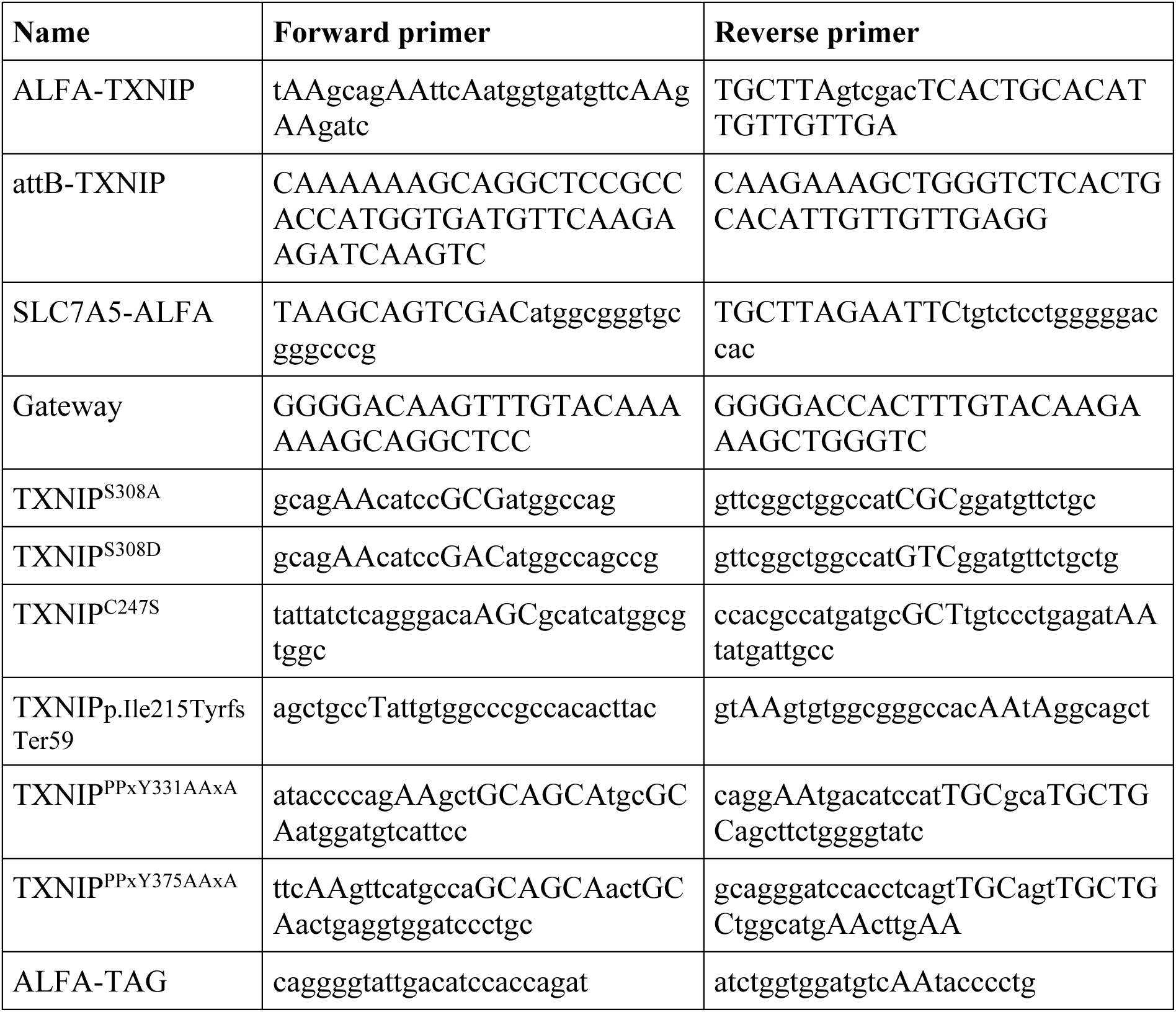
Primers for PCR-based genetic modifications and cloning.

**Supplementary Table 5:**
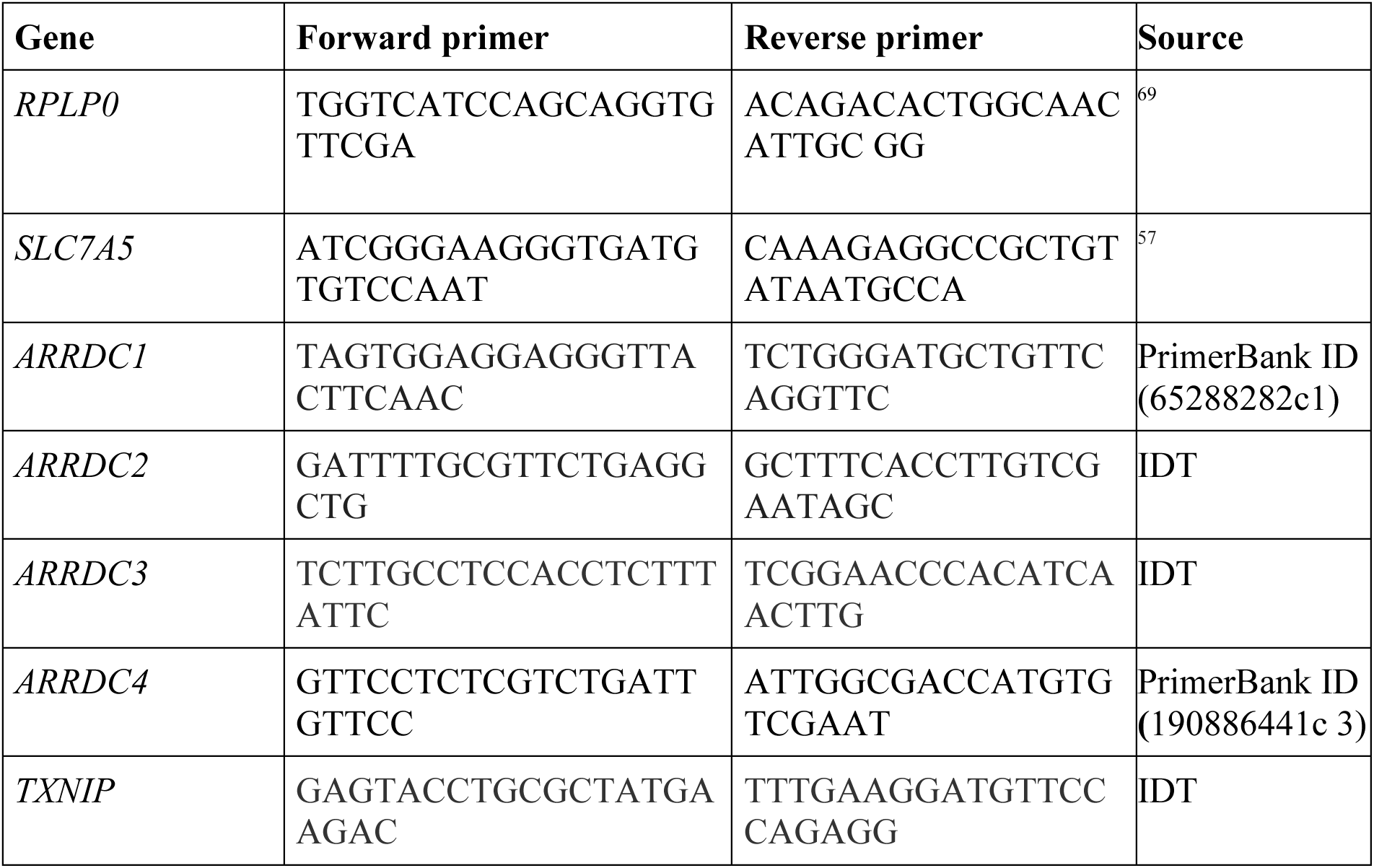
qPCR Primers used in this study.

